# The Physiological Component of the BOLD Signal: Impact of Age and Heart Rate Variability Biofeedback Training

**DOI:** 10.1101/2025.04.04.647252

**Authors:** Richard Song, Jungwon Min, Shiyu Wang, Sarah E. Goodale, Kimberly Rogge-Obando, Ruoqi Yang, Hyun Joo Yoo, Kaoru Nashiro, Jingyuan E. Chen, Mara Mather, Catie Chang

## Abstract

Aging is associated with declines in autonomic nervous system (ANS) function, including reduced heart rate variability (HRV), impaired neurovascular coupling, and diminished cerebrovascular responsiveness—factors that may contribute to cognitive decline and neurodegenerative diseases. Understanding how aging alters physiological signal integration in the brain is crucial for identifying potential interventions to promote brain health. This study examines age-related differences in how cardiac and respiratory fluctuations influence the blood oxygenation level-dependent (BOLD) signal, using two independent resting-state fMRI datasets with concurrent physiological recordings from younger and older adults. Our findings reveal significant age-related reductions in the percent variance of the BOLD signal explained by heart rate (HR), respiratory variation (RV), and end-tidal CO_2_, particularly in regions involved in autonomic regulation, including the orbitofrontal cortex, anterior cingulate cortex, insula, basal ganglia, and white matter. Cross-correlation analysis also revealed that younger adults exhibited stronger HR-BOLD coupling in white matter, as well as a more rapid BOLD response to RV and CO_2_ in gray matter. Additionally, we investigated the effects of heart rate variability biofeedback (HRV-BF) training, a non-invasive intervention designed to modulate heart rate oscillations. The intervention altered physiological-BOLD coupling in an age- and training-dependent manner: older adults who underwent HRV-BF to enhance HR oscillations exhibited a shift toward younger-like HR-BOLD coupling patterns, while younger adults who trained to suppress HR oscillations showed increased CO_2_-BOLD coupling. These findings suggest that HRV-BF may help mitigate age-related declines in autonomic or cerebrovascular function. Overall, this study underscores the role of physiological dynamics in brain aging and highlights the importance of considering autonomic function when interpreting BOLD signals. By demonstrating that HRV-BF can modulate physiological-BOLD interactions, our findings suggest a potential pathway for enhancing cerebrovascular function and preserving brain health across the lifespan.

## Section 1: Introduction

The autonomic nervous system (ANS) is a sub-branch of the peripheral nervous system that is responsible for regulating physiological functions of the body, including heart rate and respiration. It is well documented, however, that ANS health declines with age (Mather, 2024; Olivieri et al., 2024; Takla et al., 2023; Thayer et al., 2010). For example, aging is associated with decreased heart rate variability (HRV), which reflects the natural ability of the brain to modulate oscillations in heart rate (Britton et al., 2007; Jandackova et al., 2016; Mather, 2024; Reardon & Malik, 1996; Thayer et al., 2010; Zulfiqar et al., 2010). As the ANS controls cardiovascular responses, it directly influences cerebral blood flow and the dynamics of neurovascular coupling, which are vital for maintaining a healthy level of blood flow to the brain to support cognitive functions (Goadsby, 2013; Koep et al., 2022; Mankoo et al., 2023). With aging, the decline in ANS health can compromise the ability of the brain to regulate cerebral blood flow efficiently (Lu et al., 2011). This diminished neural-vascular regulation, exacerbated by age-related structural changes in the vasculature, has been linked to cognitive decline and diseases like Alzheimer’s and stroke (Han et al., 2021; Mankoo et al., 2023; Sweeney et al., 2019).

Functional magnetic resonance imaging (fMRI) is a non-invasive technique for measuring brain activity by detecting changes in local blood oxygenation levels. Since fMRI relies on the hemodynamic response, the blood oxygenation level-dependent (BOLD) signal is affected by low-frequency fluctuations in peripheral physiological processes, including natural variations in respiratory variation (RV) and heart rate (HR) (Murphy et al., 2013; Wise et al., 2004). In fMRI studies, it is common to regress out RV and HR from the BOLD signal, as these effects introduce non-neuronal variations that may confound inferences of neural activity. However, several new avenues of research have indicated that the physiological component of the fMRI BOLD signal may provide valuable information related to ANS health (Bright et al., 2020; Donahue et al., 2016; Makedonov et al., 2013; Mather & Thayer, 2018; Yang et al., 2022). For instance, areas of the brain associated with regulating naturalistic breathing rhythms were found to have high amounts of BOLD-RV coupling, and regions demonstrating high BOLD-HR coupling were found in regions of high vessel density (Chen et al., 2020), highlighting the potential for using the physiological component of the BOLD signal as an indicator of ANS health.

Due to the deterioration of ANS health with age, the effect of age on physiological-BOLD coupling is of interest. The joint dynamics of physiological signals (e.g., HR and RV) and the BOLD signal may shed light on age-related changes in autonomic and cerebrovascular health, potentially uncovering early indicators of neurodegenerative or cardiovascular diseases. One study demonstrated that aging was associated with lower levels of resting state BOLD variability and that cardiovascular health moderated this effect (Tsvetanov et al., 2021). Another study demonstrated that the hemodynamic response was smaller and slower in older adults compared to younger adults when performing audio/visual sensorimotor tasks (West et al., 2019). Further, animal models have revealed that aging may reduce neurovascular coupling due to structural changes in neurovasculature, such as vascular rarefaction and endothelial dysfunction, suggesting that the relationship between BOLD and physiological signals may be an indicator of aging (Yabluchanskiy et al., 2021). However, it is presently unclear how the spatiotemporal association of BOLD fMRI signals with low-frequency breathing and heart rate variability changes with aging.

Here, we identify age-related differences in the propagation of spontaneous heart rate and respiratory fluctuations into resting-state BOLD signals. Differences between older and younger adults prominently included brain regions that have been implicated in autonomic regulation.

Additionally, we demonstrate that HRV biofeedback training, a non-invasive paced breathing technique to modulate HR oscillations, alters the dynamics of BOLD-physiological coupling in older adults to resemble patterns typical of younger adults. Overall, this research aims to advance our understanding of how physiological signals manifest within the spatial and temporal patterns of the BOLD response, while also investigating how both the aging process and HRV biofeedback training may modulate these underlying dynamics.

## Section 2: Methods

### Section 2.1: Datasets

We included resting-state fMRI scans with high quality physiological data from 399 participants in the Nathan Kline Institute (NKI) Rockland sample (Nooner et al., 2012). Data were examined from a younger (range: 19-36, mean: 25.98, std: 4.72, n = 144) and an older (range: 50-85, mean: 63.00, std: 8.28, n = 255) group. The NKI institutional review board approved data collection, and participants gave informed consent. An MPRAGE sequence was used to retrieve an anatomical image for each subject (TR/TE = 1900/2.52 ms, FA = 9°, thickness = 1.0 mm, slices = 192, matrix = 256 × 256, FOV = 250 mm). Resting state scans were acquired using an EPI sequence (TR = 1400 ms, duration 10 min, voxel size = 2.0 mm, isotropic, FA = 65°, FOV = 224 mm). During the scan, physiological data sampled at 62.5 Hz was also collected from participants. Cardiac data was collected via photoplethysmogram (PPG) and respiration was measured using a respiration belt.

We also included resting-state fMRI scans with high quality physiological data from 110 participants in the Heart Rate Variability and Emotional Regulation (HRV-ER) dataset (Yoo et al., 2023), who were either in a younger (range: 18-30, mean: 22.19, std: 2.86, n = 59) or an older (range: 55-80, mean: 64.69, std: 6.36, n = 51) group. Resting state fMRI was acquired using multi-echo-planar imaging sequence with TR = 2.4 seconds, TE 18/35/53 ms, slice thickness = 3.0 mm, flip angle = 75°, field of view = 240 mm, voxel size = 3.0 × 3.0 × 3.0 mm, 175 volumes, and acquisition time = 7 min. In-scan cardiac data was collected using PPG, CO_2_ levels were collected using capnography, and respiration was measured using a breathing belt. Physiological data was initially collected at 10 kHz but was downsampled to 1 kHz. Scans were collected from all participants both before and after a 5 week HRV biofeedback training intervention.

During the 5-week HRV biofeedback training, half of the participants completed daily sessions in which they did slow paced breathing while receiving biofeedback to increase their heart rate oscillations (Osc+), while the other half received biofeedback to decrease heart rate oscillations (Osc−). In weekly lab sessions, participants in the Osc+ condition practiced breathing at different paces to identify their resonance frequency, aiming to maximize heart rate oscillations using biofeedback from the emWave Pro software. In the Osc− condition, participants used their own strategies to reduce heart rate oscillations, receiving feedback via a <calmness= score, which increased as their heart rate oscillations decreased. In between the weekly lab sessions, training was performed twice daily at home for 20 minutes each using the breathing pace (Osc+) or strategy (Osc-) identified as most effective for increasing (Osc+) or reducing (Osc-) heart rate oscillations. Of the 110 participants for which baseline resting-state fMRI scans and in-scanner physiological recordings were obtained, 78 participants (20 younger Osc+, 21 younger Osc-, 19 older Osc+, 18 older Osc-) also had the fMRI scans and physiological recordings data after the 5-week HRV biofeedback training.

### Section 2.2: Imaging Data Preprocessing

For anatomical data in both NKI and HRV-ER datasets, T1-weighted images were skull-stripped using FSL’s Brain Extraction Tool (BET), followed by intensity non-uniformity correction with AFNI’s *3dUnifize*.

In the NKI dataset, motion correction of the fMRI data was carried out using FSL *mcflirt*. ICA FIX (Griffanti et al., 2014) was additionally used to further correct for motion artifacts including high-frequency physiologically-coupled motion artifacts and other MRI acquisition-related artifacts.

ICA FIX functions as an independent component classifier, and was trained using data from 25 subjects with a balanced age distribution from the NKI dataset. For training, each independent component was manually labeled as either “noise” or “not noise,” enabling ICA FIX to identify and remove noise components in the remainder of the dataset.

In the HRV-ER dataset, fMRI data underwent motion correction using AFNI’s *3dvolreg*, applied to the second echo, with the resulting motion parameters used to align the first and third echoes via *3dAllineate*. Slice timing correction was performed with AFNI’s *3dTshift*. Multi-echo fMRI data were denoised using *tedana*, which applied multi-echo independent component analysis (ME-ICA) to separate BOLD from non-BOLD signals (Kundu et al., 2012, 2017).

In both datasets, spatial normalization of functional data to MNI space was achieved by first aligning functional volumes to the T1-weighted anatomical images with FSL’s *epi_reg*, followed by ANTs-based registration to the MNI152 2mm template. The ANTs routine *antsApplyTransforms* was used to apply this nonlinear transformation to the T1-registered functional data. Smoothed functional images (3mm FWHM) were generated using AFNI’s *3dmerge*. Nuisance regression, including fourth-order polynomial detrending, was performed with *3dDetrend*, followed by the addition of the mean signal using *3dTstat*.

### Section 2.3: Physiological Data Preprocessing

Physiological data, including heart rate (HR), respiratory variation (RV), and end-tidal CO_2_, were processed to create fMRI regressors aligned with the fMRI time series. To calculate HR, the raw photoplethysmography (PPG) signal was bandpass-filtered between 0.5 and 2 Hz using a second-order Butterworth filter to enhance heartbeat clarity for peak detection. Peaks were identified using a minimum peak height threshold of 5% of the interquartile range, and inter-beat intervals (IBI) were calculated. After visual inspection of the IBI time series for artifacts (e.g., due to poor peak detection), we interpolated over any instances where artifacts occurred. HR was then calculated as the inverse of the median IBI per minute using 6-second sliding windows centered at each fMRI TR. Metrics of heart rate variability (HRV), such as RMSSD, high frequency (HF), and low frequency (LF) power, were also derived from the IBI. The HF power band was defined between 0.15 – 0.4 Hz, and the LF power band was defined between 0.04 – 0.15 Hz (Burr, 2007).

RV was calculated as the temporal standard deviation of the raw respiration waveform within 6-second windows centered at each TR. To account for individual differences in torso size, the RV signal was normalized between -1 and 1 for each participant. RV normalization was performed by first determining lower and upper bounds at the 1.45th and 98.55th percentiles using a histogram-based approach. The mean RV within these bounds was then subtracted to center the data approximately around zero, and this mean was also subtracted from the lower and upper bounds. Finally, positive and ngeative values were divided by their respective adjusted upper or lower bounds, yielding a normalized range approximately spanning -1 to 1.

The HRV-ER dataset also included capnography recordings for measuring CO_2_. To account for time delays in the capnography signal, it was time-shifted to maximize its alignment with the respiration waveform, reflecting the expected inverse correlation between the two signals. End-tidal CO_2_ values were then extracted from the re-aligned capnography peaks and resampled to match the fMRI TR. For both datasets, breathing rate was also calculated in breaths per minute using the *NeuroKit2* package in Python (Makowski et al., 2021).

### Section 2.4: Modeling relationships between BOLD and Physiological Signals

The physiological data were used to create regressors for modeling the impact of resting-state physiological fluctuations on BOLD signal variance. To account for respiratory effects, we convolved the RV signal with the primary Respiratory Response Function (RRF) (Birn et al., 2008), as well as temporal and dispersive derivative basis functions of the RRF. This allowed us to model regional differences in the propagation of RV effects on the BOLD signal (Chen et al., 2020). Similarly, heart rate (HR) was convolved with the primary Cardiac Response Function (CRF) (Chang et al., 2009) and its temporal and spatial derivative basis functions. The RRF and CRF model the physiological impulse responses, capturing how neural and vascular changes due to physiological fluctuations influence the BOLD signal. A schematic for this model is shown in *Figure 1*.

**Figure 1.**
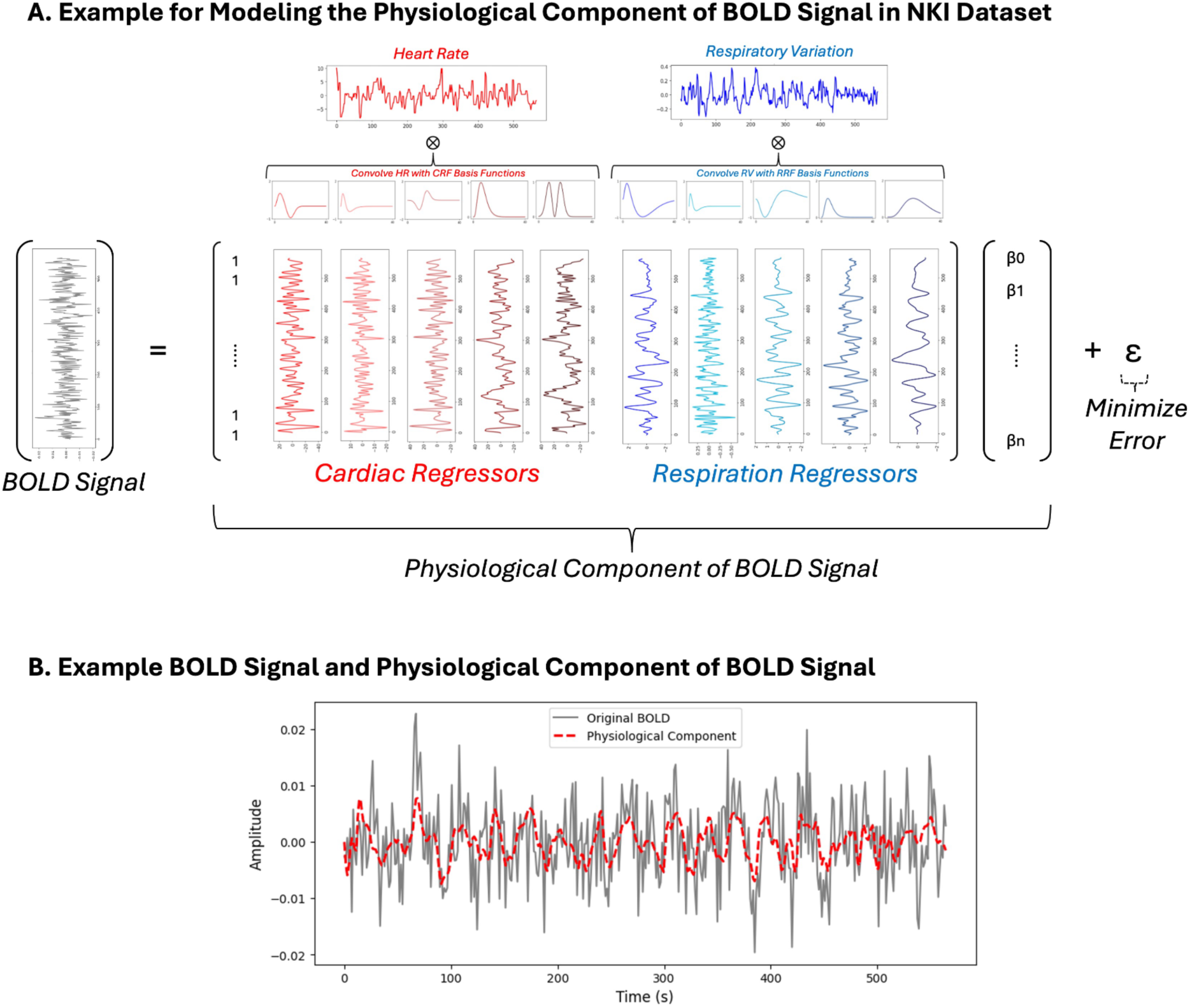
Schematic for the model to determine the physiological component of the BOLD signal. A) After detrending and normalizing heart rate and respiratory variation, the signals are convolved with CRF and RRF basis functions. For every voxel, a general linear model is used to find beta weights for each of the cardiac and respiration regressors to minimize the error from the original BOLD signal. B) Example of an original BOLD signal (normalized to percent signal change) and the corresponding physiological component.

In the HRV-ER dataset, we also modeled the impact of end-tidal CO_2_ on the BOLD signal by convolving it with a previously parameterized double-gamma end-tidal CO_2_ response function (Golestani et al. 2015), and its temporal derivative. The Golestani et al. (2015) basis functions were chosen to model end-tidal CO_2_ response over other commonly used response functions for modeling the resting-state BOLD response to CO_2_, such as the canonical hemodynamic response function (Yao et al. 2021), due its ability to better capture the delayed response of the BOLD signal in older adults to CO_2_ compared to younger adults, which is detailed more in Section 3.3 and *Supplemental Figure 1*.

For each dataset, we fitted three separate linear models to assess the effect of physiological activity on the BOLD signal: (1) a model with both cardiac and respiration regressors, (2) a model with only cardiac regressors, and (3) a model with only respiration regressors. These models provided the BOLD signal variance explained by both physiological factors combined, cardiac activity alone, and respiration alone, respectively, for each voxel across all participants. Since the signal quality for respiration belt data collected in the HRV-ER dataset was deemed poor after visual inspection, we used end-tidal CO_2_ as the measure of respiration in this dataset. In the NKI dataset, RV was used as the measure of respiration. Both datasets used heart rate as the measure of cardiac activity. To determine the relative contribution of cardiac and respiratory activity to the BOLD signal, we calculated the percent variance explained (PVE) by dividing the variance in the BOLD signal explained by each model by the total variance in the original BOLD signal for every voxel.

In addition to the PVE analysis, whole-brain voxel-wise cross-correlations were computed to examine the temporal relationship between physiological signals and BOLD fluctuations. Cross-correlations were calculated between the BOLD signal and HR/RV for the NKI dataset and between the BOLD signal and HR/end-tidal CO_2_ for the HRV-ER dataset. These correlations were computed across time lags ranging from -2.8s to 21.4s in the NKI dataset, with 1.4s increments, and from -2.4s to 21.6s in the HRV-ER dataset, with 2.4s increments, corresponding to the respective TRs of each dataset.

To further capture global trends at a macro level, cross-correlations were also computed using the BOLD signal averaged across three major tissue compartments: gray matter, white matter, and ventricles. This approach allowed for the examination of broader patterns in physiological signal propagation beyond voxel-level resolution. Statistical comparisons were conducted at each time lag using t-tests, with multiple comparisons corrected using Bonferroni correction.

### Section 2.5: Whole-Brain Statistical Testing

Whole-brain maps of physiological signal propagation into BOLD fMRI data were compared in older versus younger adults using the HRV-ER and NKI datasets, focusing on the baseline (pre-intervention) condition in the HRV-ER dataset. Statistical differences in percent variance explained maps, or voxel-wise cross-correlation maps, were assessed using two-sample t-tests. To evaluate the effects of the HRV biofeedback intervention, pre- and post-intervention data from the subset of HRV-ER participants with both conditions were used to compute post-minus-pre differences separately for younger and older adults within each intervention condition (Osc+ and Osc-). To assess the effect of age on the intervention response, these post-pre difference maps were then compared between age groups using two-sample t-tests. For all statistical analyses of whole-brain maps, multiple comparisons were corrected using the Threshold-Free Cluster Enhancement (TFCE) algorithm with 5000 permutations.

## Section 3: Results

### Section 3.1: Physiological Differences by Age

Information about physiological metrics in the participants of the two datasets can be found in Table 1 and Table 2, and are summarized in *Figure 2*. When calculating summary statistics, we identified outliers using the interquartile range (IQR) method, where values falling below Q1-1.5×IQR or above Q3+1.5×IQR were excluded from analysis for that specific metric only, while retaining the participant’s valid data for other measurements. Since RMSSD, low frequency (LF) HRV, and high frequency (HF) HRV are not normally distributed, they were transformed using the natural logarithm before statistical analysis. As detailed in Table 1 and Table 2, the older adults exhibited significantly lower ln(LF HRV) and ln(HF HRV) than younger adults. Although older adults had lower RMSSD than younger adults, this difference was not significant in the NKI dataset, although it was in the HRV-ER dataset. As shown in Table 1, older adults also exhibited higher standard deviation in RV than younger adults. However in both datasets, average HR was not significantly different between the two age groups. In addition, breathing rate was significantly higher in younger adults in both datasets.

**Table 1:** Physiological differences by age in NKI dataset. Mean and variance for all measures are shown, along with t-test statistics comparing averages across age groups.

| Metric | Younger Adults<br>(n = 144) | Older Adults<br>(n = 255) | t-statistic | p-value |
| --- | --- | --- | --- | --- |
| ln RMSSD | 4.197 (0.410) | 4.186 (0.640) | 0.144 | 0.885 |
| ln LF | 6.879 (1.239) | 6.373 (2.211) | 3.553**** | 4.266e-04 |
| ln HF | 7.177 (1.661) | 6.814 (2.608) | 2.310* | 2.142e-02 |
| Avg HR | 69.046 (82.872) | 67.442 (80.538) | 1.696 | 9.062e-02 |
| SD RV | 0.094 (0.001) | 0.104 (0.002) | -2.551* | 1.115e-02 |
| Breathing Rate | 17.700 (9.253) | 15.610 (11.803) | 5.922**** | 7.212e-09 |

\* p < 0.05, \*\* p < 0.01, \*\*\* p < 0.005, \*\*\*\* p < 0.001

**Table 2:** Physiological differences in HRV-ER dataset by age and Osc condition. Mean and variance for all measures are shown, along with t-test statistics comparing averages across age groups and between pre– and post–intervention within each group.

| Metric | Younger Osc+<br>(n = 20) |  |  | Young Osc-<br>(n = 21) |  |  | Old Osc+<br>(n = 19) |  |  | Old Osc-<br>(n = 18) |  |  | Young vs Old<br>Pre t-stats |
| --- | --- | --- | --- | --- | --- | --- | --- | --- | --- | --- | --- | --- | --- |
|  | Pre | Post | t-stat | Pre | Post | t-stat | Pre | Post | t-stat | Pre | Post | t-stat | t-stat |
| ln RMSSD | 3.89<br>(0.36) | 3.98<br>(0.36) | -0.985 | 4.06<br>(0.31) | 4.05<br>(0.22) | 0.049 | 3.13<br>(0.39) | 3.53<br>(0.52) | -1.397 | 3.42<br>(0.72) | 3.14<br>(0.68) | 1.105 | 5.140**** |
| ln LF | 6.53<br>(0.78) | 6.73<br>(0.72) | -0.938 | 6.30<br>(0.75) | 6.49<br>(1.11) | -0.902 | 5.30<br>(1.65) | 6.01<br>(2.30) | -1.699 | 5.82<br>(1.79) | 5.56<br>(2.64) | 0.609 | 4.301**** |
| ln HF | 6.71<br>(1.20) | 7.00<br>(1.43) | -1.283 | 7.04<br>(1.26) | 7.02<br>(0.87) | 0.072 | 4.82<br>(1.34) | 5.87<br>(1.64) | -2.371* | 5.78<br>(2.44) | 5.49<br>(2.48) | 0.553 | 6.495**** |
| Avg HR | 67.64<br>(102.85) | 66.65<br>(79.07) | 0.561 | 66.93<br>(82.67) | 65.91<br>(63.38) | 0.589 | 65.68<br>(39.37) | 62.90<br>(65.45) | 2.233* | 69.00<br>(105.87) | 68.04<br>(100.84) | 0.721 | -0.153 |
| Avg CO<sub>2</sub> | 38.48<br>(25.69) | 39.30<br>(13.53) | -0.574 | 39.53<br>(22.44) | 39.10<br>(35.22) | 0.290 | 39.74<br>(14.10) | 39.30<br>(18.29) | 0.607 | 40.68<br>(12.82) | 42.08<br>(16.28) | -2.753* | -0.019 |
| Breathing<br>Rate | 16.90<br>(16.08) | 15.50<br>(19.07) | 1.744 | 18.03<br>(7.60) | 18.13<br>(7.53) | -0.212 | 15.87<br>(11.98) | 14.86<br>(17.80) | 1.586 | 13.73<br>(13.59) | 13.66<br>(10.36) | 0.104 | 2.767** |

\* p<0.05, \*\* p<0.01, \*\*\*p<0.005, \*\*\*\*p<0.001

**Figure 2:**
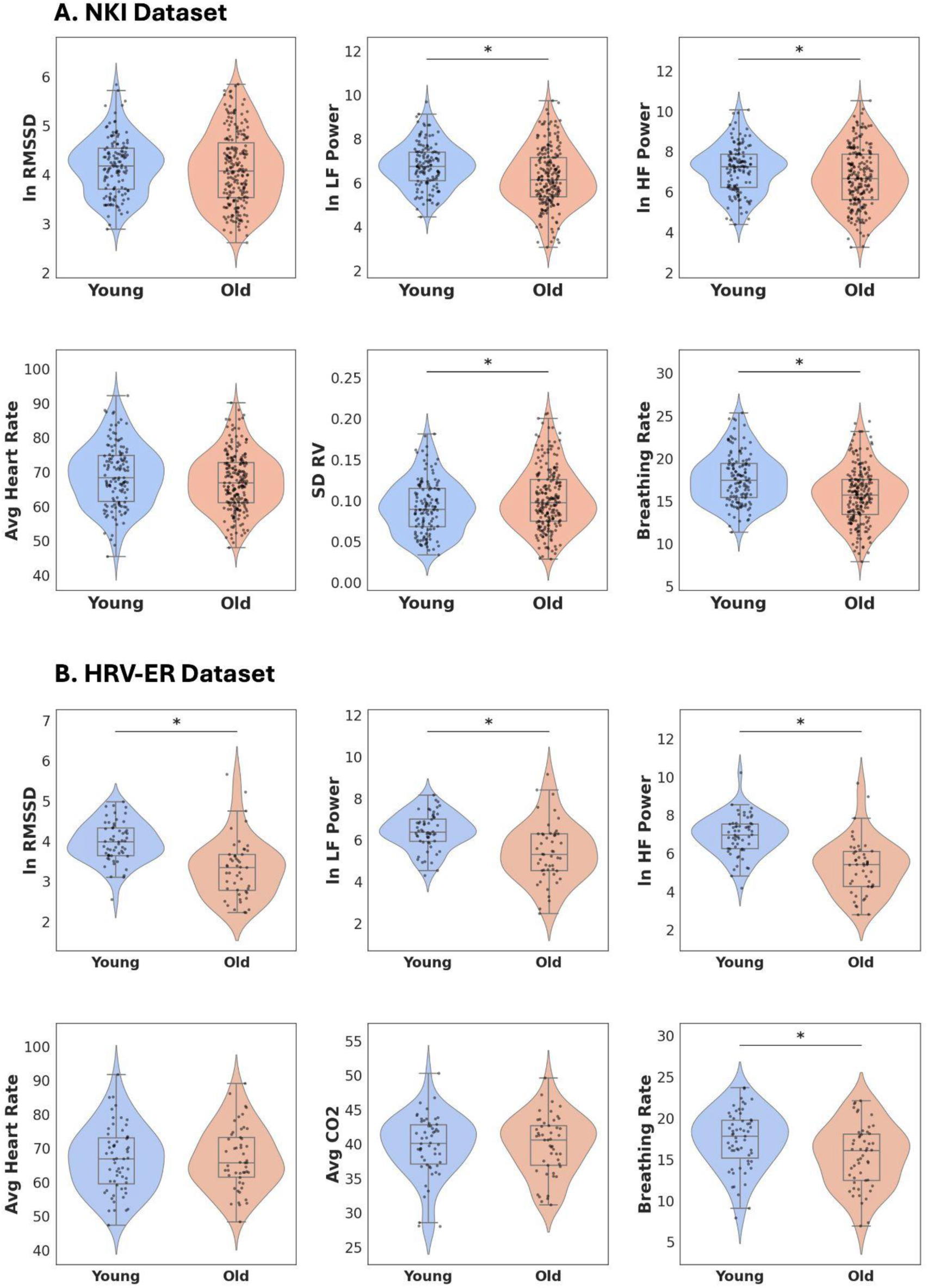
Log-transformed metrics of HRV (RMSSD, LF Power, HF Power) and low-frequency peripheral physiology across the A) NKI dataset and B) HRV-ER dataset. Standard box and whisker plots are overlaid over the violin plots. *p < 0.05

### Section 3.2: Age Differences in the Percent Variance of the BOLD Signal Explained by Peripheral Physiological Measures

To examine the extent to which the fMRI BOLD signal was associated with peripheral physiological measures, we convolved heart rate and respiration time courses with their respective physiological impulse response basis functions to obtain cardiac and respiration regressors, which we then fit to the BOLD signal in each voxel using a general linear model (Section 2.4). For the comparison of age-related differences in the HRV-ER dataset, subjects’ percent variance explained maps were calculated based only on their baseline scan before the HRV biofeedback intervention.

Results are presented in Figure 3. In both datasets, the percent variance in BOLD signal explained by heart rate and respiration was significantly greater in younger adults, compared to older adults, in the orbital frontal cortex (OFC), lateral ventricle, basal ganglia, and white matter (*Figure 3A*). In the NKI dataset, percent variance of BOLD explained by heart rate and respiration was also significantly higher in the ACC and insula; these were also amongst the regions showing the strongest mean age-related differences in the HRV-ER dataset, though they did not survive the statistical threshold. The maps of percent variance of BOLD explained by heart rate (*Figure 3B*) closely resembled the statistical maps resulting from jointly fitting heart rate and respiration (*Figure 3A*). In both datasets, percent variance of BOLD explained by heart rate was higher in younger adults than older adults in the lateral ventricles, basal ganglia, and white matter (*Figure 3B*). There were also subthreshold differences in the HRV-ER dataset that were significant in the NKI dataset in the ACC, insula, and OFC. Interestingly, the percent variance in BOLD explained by respiration (e.g. RV in the NKI dataset and CO_2_ in the HRV-ER dataset) showed different age-related differences. In the NKI dataset, BOLD variance accounted for by RV was significantly higher in younger adults in OFC, insula, lateral ventricles, and ACC (*Figure 3C*). In the HRV-ER dataset, older adults had slightly higher percent variance in BOLD accounted for by CO_2_ than younger adults in white matter and occipital cortex when comparing the group average maps; however, these differences were not statistically significant.

**Figure 3.**
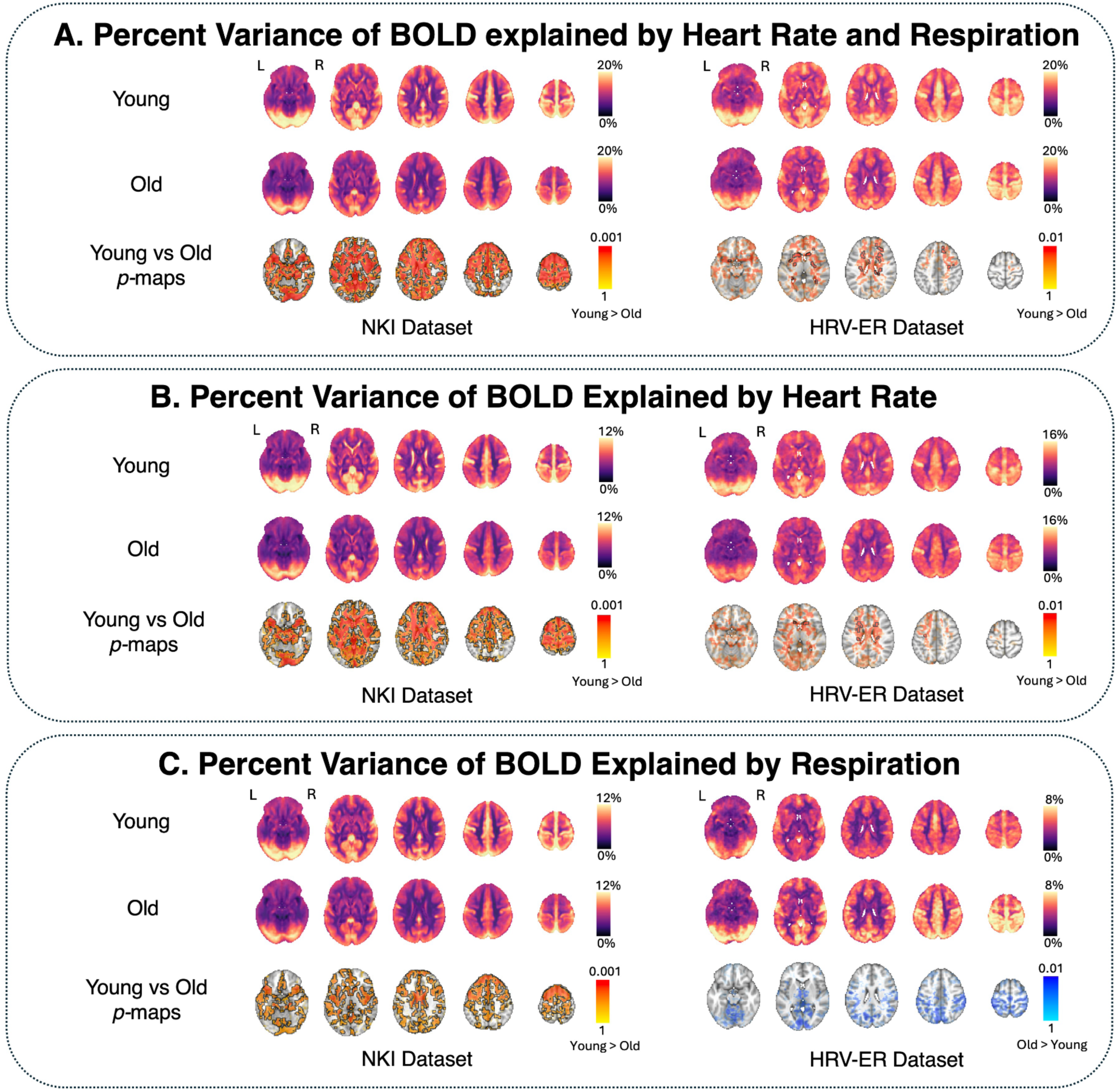
Percent variance of BOLD signal explained by low-frequency peripheral physiology across all voxels. Three separate models were used to determine the variance in the BOLD signal explained by A) both heart rate and respiration, B) only heart rate, and C) only respiration. In each panel, age group averages are shown for young and old participants. Voxels in which percent variance explained in younger adults was statistically significantly greater than older adults (*p* < 0.05 TFCE-corrected) are outlined in black, and alpha-fading was used to highlight sub-threshold voxels. For visualization purposes in the NKI dataset, *p* values were transformed using the natural logarithm to improve the contrast between highly significant voxels (e.g. *p* < 0.01). The brain slices shown are at z = - 16 mm, 4 mm, 24 mm, 44 mm, and 64 mm in standard MNI152 space.

### Section 3.3: Age Differences in the Cross-Correlation Between the BOLD Signal and Physiological Measures at Different Time Lags

To further examine how age impacts the dynamic association between physiological and BOLD signals, we computed the cross-correlations between the two signals (Section 2.5). In addition to revealing information about the temporal dynamics of BOLD-physiological coupling, a cross-correlation analysis does not assume specific hemodynamic impulse response models in the relationship between physiological and BOLD signals. Results for the NKI Dataset are shown in *Figure 4* and results for the HRV-ER Dataset are shown in *Figure 5*, with significant clusters outlined in the figures (and discussed below) at TFCE-corrected thresholds of *p*<0.05.

**Figure 4.**
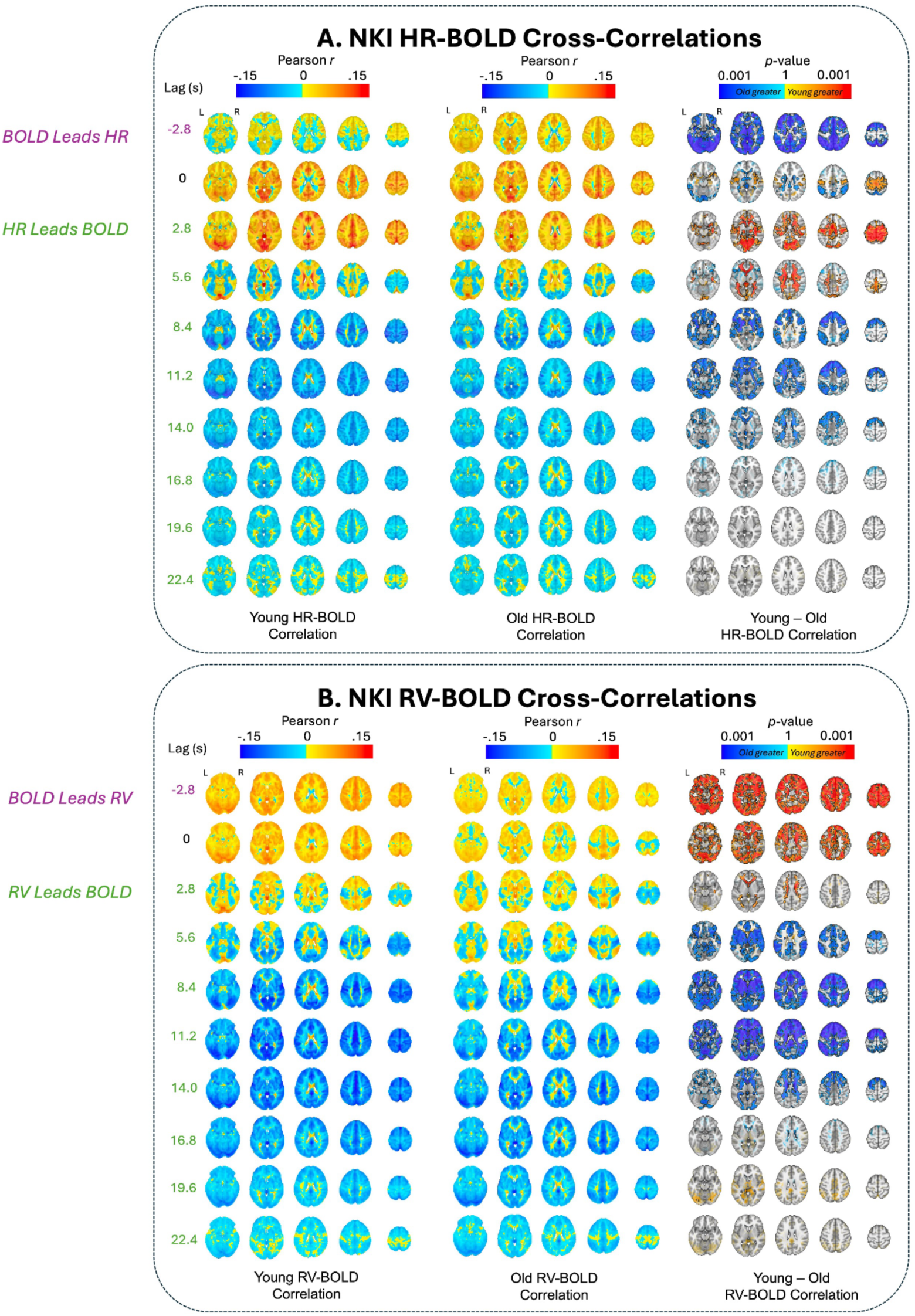
A) HR-BOLD and B) RV-BOLD cross-correlations for different time lags in the NKI dataset. Negative lags indicate that BOLD leads the physiological measure, and vice versa for positive lags. Age group averages for Pearson *r* coefficients are plotted at each lag. Significant voxels by age group at *p* < 0.05 (TFCE-corrected) are also outlined in black at each lag, along with alpha fading to show sub-threshold voxels. Red voxels indicate that young adult *r* values are greater than old adults, and blue voxels indicate that old adult *r* values are more positive (or less negative) than young adults. For visualization purposes, *p* values were transformed using the natural logarithm to improve the contrast between highly significant voxels (e.g. *p* < 0.01). The brain slices shown are at z = -16 mm, 4 mm, 24 mm, 44 mm, and 64 mm in standard MNI152 space.

**Figure 5.**
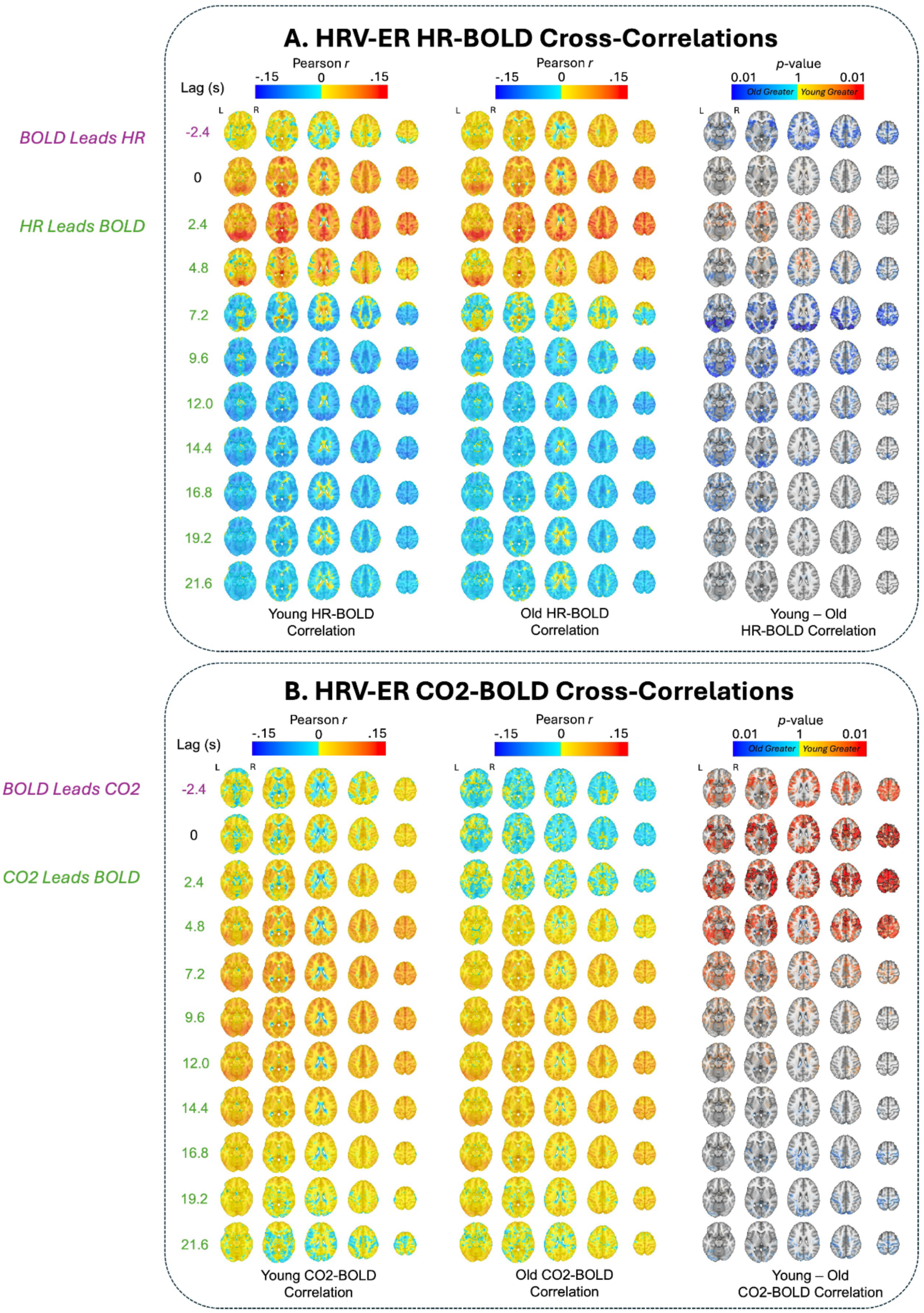
A) HR-BOLD and B) CO_2_-BOLD cross-correlations for different time lags in the HRV-ER dataset at baseline. Negative lags indicate that BOLD leads the physiological measure, and vice versa for positive lags. Age group averages for Pearson *r* coefficients are plotted at each lag. Significant voxels by age group at *p* < 0.05 (TFCE-corrected) are also outlined in black at each lag, along with alpha fading to show sub-threshold voxels. Red voxels indicate that young adult *r* values are greater than old adults, and blue voxels indicate that old adult *r* values are greater than young adults. The brain slices shown are at z = -16 mm, 4 mm, 24 mm, 44 mm, and 64 mm in standard MNI152 space.

In the NKI dataset, the older adults had significantly higher HR-BOLD cross-correlation across much of the brain at a time lag of -2.8s. On the other hand, younger adults had significantly higher HR-BOLD cross-correlation in lags 2.8 seconds and 5.6 seconds in the ventricles, white matter, and ACC (*Figure 4A*). In addition, younger adults had stronger negative HR-BOLD cross-correlations in the prefrontal cortex (PFC), OFC, and insula in lags 8.4 seconds and 11.2 seconds, as well as in the lateral ventricles in lags 11.2 seconds and 14.0 seconds (*Figure 4A*). Similar spatiotemporal trends were present in the HRV-ER dataset, albeit with smaller and/or nonsignificant effect sizes (*Figure 5*). Here, the older adults also had significantly higher HR-BOLD cross-correlation across most of the brain at time lag of -2.4s. The younger adults had higher subthreshold HR-BOLD cross-correlation in lags 2.4 and 4.8 seconds in a small number of voxels in the white matter, ventricles, ACC, and OFC (*Figure 5A)*. Younger adults also had significantly more negative HR-BOLD cross-correlation in occipital cortex at lag 7.2 seconds (*Figure 5A)*. In addition, younger adults had more negative sub-threshold differences in HR-BOLD correlations throughout the gray matter from lags 7.2 seconds to 19.2 seconds (*Figure 5A)*. These results are further supported by *Figure 6A*, which show that in the NKI dataset, HR-BOLD cross correlations are significantly higher in younger adults at earlier lags (between 3 and 6 seconds) in white matter and ventricles, and significantly more negative in younger adults at later lags (between 10 -15 seconds) across the whole brain. The HRV-ER dataset shows similar spatiotemporal trends with smaller and/or nonsignificant effect sizes (*Figure 6A*).

**Figure 6.**
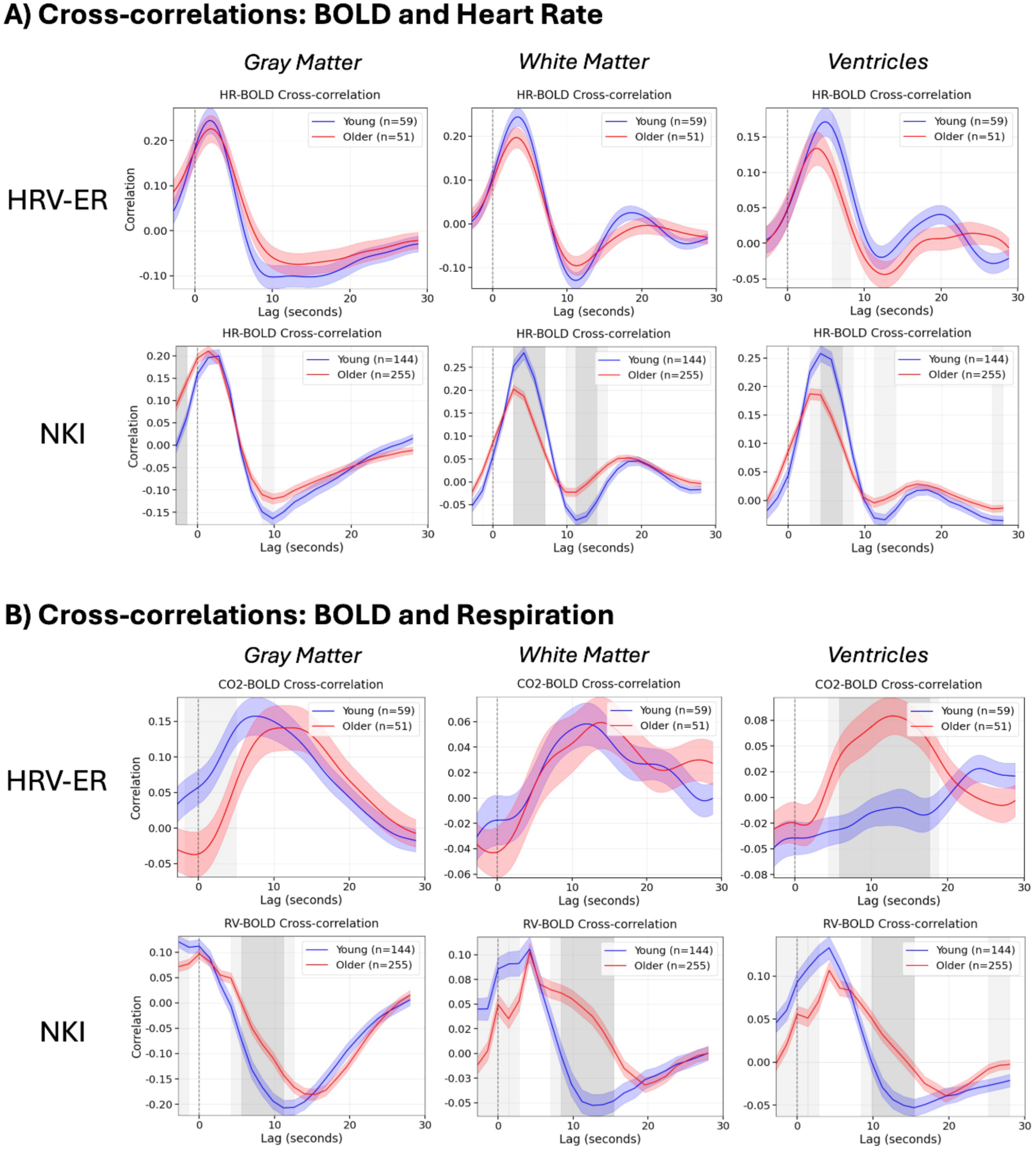
Cross-correlations between the BOLD signal and A) heart rate and B) respiration (i.e., CO_2_ for HRV-ER, RV for NKI), averaged across three tissue types: gray matter, white matter, and ventricles. In the HRV-ER data, both BOLD signal and CO_2_/HR were upsampled to TR = 0.2 seconds before cross-correlation calculation, but no upsampling occurred in the NKI data. Group averages for old and young adults are plotted along with shading for standard error. Lags where the cross correlations between older and younger adults were statistically significant (*p*<0.05) after Bonferroni correction are plotted in dark gray, and lags where the cross correlation passed a *p*<0.05 uncorrected threshold are plotted in light gray.

In the NKI dataset, RV-BOLD cross-correlations were significantly higher in younger adults in the ventricles at lags -2.8 seconds, 0 seconds, and 2.8 seconds, as well as across the gray matter at lags -2.8 seconds and 0 seconds (*Figure 4B*). From lags 5.6 seconds to 14 seconds, younger adults exhibited significantly more negative RV-BOLD cross-correlations in the ACC, basal ganglia, and PFC at lags 5.6, 8.4, and 11.2 seconds, with additional negative RV-BOLD correlations observed in the lateral ventricles at lags 8.4, 11.2, and 14 seconds (*Figure 4B*). Similarly, in the HRV-ER dataset, younger adults exhibited significantly higher (*p* < 0.05 TFCE-corrected) CO_2_-BOLD cross-correlations in the OFC, insula, and ACC at lags 0, 2.4, and 4.8 seconds, with subthreshold differences appearing at lag -2.4 seconds and between 7.2 and 12 seconds (*Figure 5B*). *Figure 6B* further illustrates this pattern, showing that in the NKI dataset, RV-BOLD cross-correlations were higher (*p < 0.05* uncorrected) in younger adults compared to older adults between 0 and 3 seconds in the white matter and ventricles. Additionally, younger adults exhibited significantly more negative RV-BOLD cross-correlations in all three tissue types at later lags between 7 and 12 seconds. This trend is consistent with the HRV-ER dataset, where CO_2_-BOLD cross-correlations in younger adults were greater in gray matter between -2 and 5 seconds (*p < 0.05* uncorrected). These findings highlight an earlier onset of BOLD responsiveness to respiration in younger adults, as supported by both datasets.

### Section 3.4: Age- and Condition-Dependent Modulations of HR- and CO_2_-BOLD Coupling Following HRV Biofeedback

To assess the effects of the HRV-biofeedback on physiological signal propagation into the fMRI BOLD signal, we computed whole-brain HR-BOLD and CO_2_-BOLD cross correlations both before and after 5 weeks of each intervention separately for both age groups (Section 2.5). For older adults, the HR-BOLD cross correlation was significantly more negative (*p* < 0.05 Bonferroni-corrected) after Osc+ in the gray matter from 6 to 11 seconds, and this trend was present in the gray matter from 5 to 17 seconds (*p* < 0.05 uncorrected) (*Figure 7A*). This effect is further evident in *Figure 8A*, which shows the voxelwise equivalent of this whole-brain average score. Here, from 7.8 to 16.2 seconds, older adults had more negative HR-BOLD cross correlations (subthreshold) throughout the gray matter after Osc+, and this result reached statistical significance between 9.6 and 12.0 seconds in the occipital cortex.

**Figure 7.**
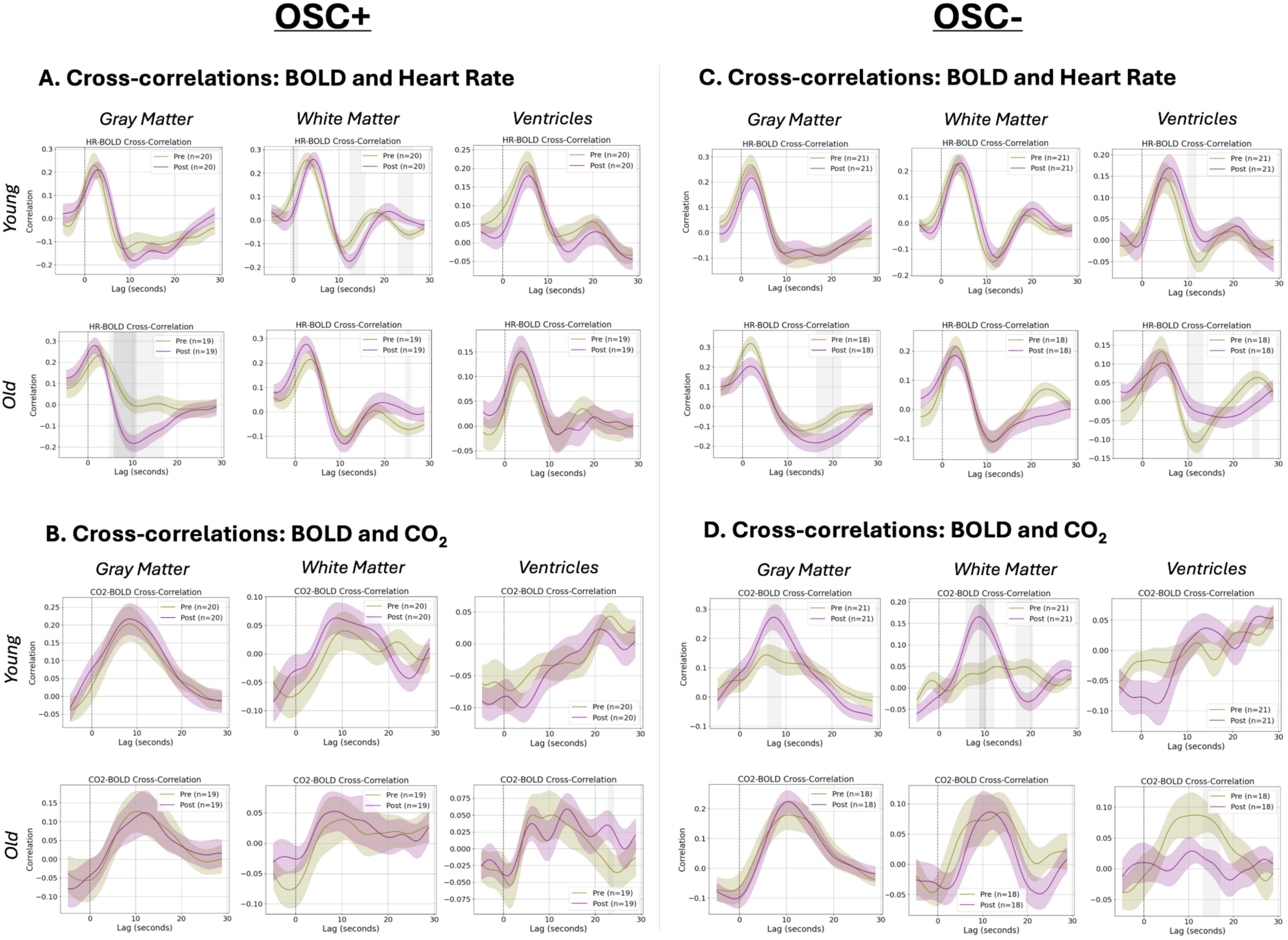
HRV biofeedback training results on BOLD-physio cross-correlations. Cross-correlations between the BOLD signal (averaged across gray matter, white matter, and ventricles) and A) HR and B) CO_2_, before and after the Osc+ condition, are plotted. Cross-correlations between the BOLD signal (averaged across gray matter, white matter, and ventricles) and C) HR and D) CO_2_, before and after the Osc-condition, are plotted. Group averages are plotted along with standard error shading at every time point. Both BOLD signal and CO_2_/HR were unsampled to TR = 0.2 seconds before cross-correlation calculation. Lags where the cross correlations between pre and post intervention were statistically significant (p<0.05) after Bonferroni correction are plotted in dark gray, and lags where the difference passed a *p*<0.05 uncorrected threshold are shown in light gray.

**Figure 8.**
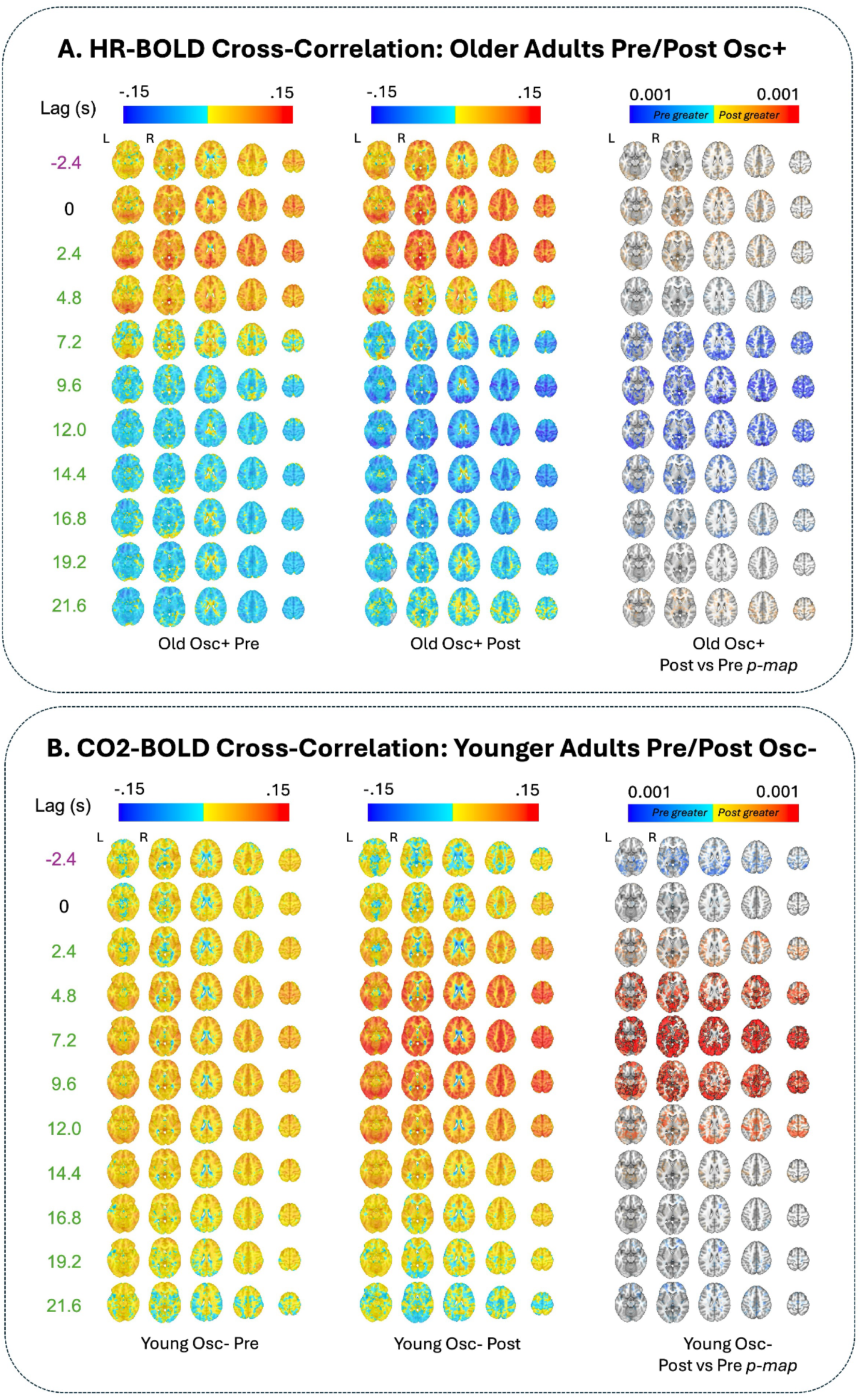
A) HR-BOLD whole-brain cross correlation plots for older adults before and after the Osc+ intervention. B) CO_2_-BOLD whole-brain cross-correlation plots for younger adults before and after the Osc-intervention. Group averages for Pearson r coefficients are plotted at each lag. Significant voxels by age group at p < 0.05 (TFCE-corrected) are also outlined in black at each lag, along with alpha fading to show sub-threshold voxels. Red voxels indicate that post Osc intervention r values are greater than pre Osc intervention r values, and vice versa for blue voxels. The brain slices shown are at z = -16 mm, 4 mm, 24 mm, 44 mm, and 64 mm in standard MNI152 space.

In addition, for younger adults, the CO_2_-BOLD cross correlation was significantly higher (*p* < 0.05 Bonferroni-corrected) post Osc-than pre Osc- in the white matter between 8 and 10 seconds, with the same trend also present in gray matter from 6 to 9 seconds and white matter from 5 to 12 seconds (*p* < 0.05 uncorrected) *(Figure 7D)*. This result is further supported by *Figure 8B,* which shows that CO_2_-BOLD cross correlation was significantly higher for younger adults after Osc+ than before at lags 4.8 seconds in the gray matter, and at lags 7.2 and 9.6 seconds in the gray and white matter. In addition, for participants in the Osc- condition, younger adults exhibited significantly greater (*p <* 0.05 TFCE-corrected) post-pre BOLD variance explained by HR and CO_2_ in the left temporal lobe (*Supplemental Figure 3A)*. This effect appears to be driven primarily by BOLD-CO_2_ coupling rather than BOLD-HR coupling, as younger adults had greater BOLD variance explained by CO_2_ in left OFC, left insula, left temporal cortex along with the lateral ventricles, bilateral occipital cortices, and bilateral PFC (*Supplemental Figure 3C*). However, most of these voxel-level differences did not reach statistical significance after correcting for multiple comparisons using TFCE with 5000 permutations.

## Section 4: Discussion

### Section 4.1: General Findings

This study examined the impact of aging, and of an HRV biofeedback intervention paradigm, on the dynamics of heart rate and respiration effects in the fMRI BOLD signal across two independent datasets. Across both datasets, we observed significant age-related differences in HRV metrics, with younger participants demonstrating higher levels of RMSSD, HF HRV, and LF HRV compared to older participants. Younger adults also exhibited higher BOLD signal variance explained by heart rate and respiration across both datasets in areas such as the white matter, OFC, ventricles, insula, and ACC, results that were also mirrored in the cross-correlation analysis. In both the CO_2_-BOLD cross-correlations in the HRV-ER and the RV-BOLD cross-correlations in the NKI dataset, younger adults exhibited significantly earlier response onset than in older adults. Finally, HRV biofeedback training was found to modulate physiological signal propagation into the BOLD signal in a condition- and age-dependent manner. Specifically, in younger adults the Osc-condition increased CO_2_-BOLD coupling, and in older adults, the Osc+ condition caused HR-BOLD coupling to resemble a pattern more typical of younger adults. These findings offer new insights into the relationship between autonomic nervous system (ANS) function and brain aging, and we discuss their implications below.

### Section 4.2: HRV Metrics and ANS Regulation

The observed reduction in HRV metrics (ln RMSSD, ln LF, ln HF) in older adults aligns with prior research linking aging to diminished ANS regulation (Jandackova et al., 2016; Reardon & Malik, 1996; Voss et al., 2012). HRV reflects the dynamic adaptability of the ANS in response to internal and external stimuli, encompassing both sympathetic and parasympathetic activity.

Lower HRV, as observed in older adults, is indicative of reduced autonomic flexibility and cardiovascular adaptability, which have been associated with impaired baroreceptor sensitivity, vascular stiffening, and altered neurocardiac signaling pathways (Yugar et al., 2023; Ziegler, 2021). Consequently, cerebral blood flow becomes more susceptible to systemic blood pressure variations, potentially causing cerebral hypoperfusion or hyperperfusion (Ogoh & Tarumi, 2019). Over time, these disruptions may contribute to structural changes in the brain, including white matter lesions and microvascular damage, which are strongly linked to cognitive decline and neurodegenerative diseases such as Alzheimer’s (Badji et al., 2019; Han et al., 2021; Reeve et al., 2024).

### Section 4.3: Age-related Differences in Physiological-BOLD Dynamics

To model the contribution of physiological signals to BOLD variance, we convolved basis functions for the cardiac response function (CRF) with HR (Chang et al., 2009; Chen et al., 2020) and for the respiratory response function (RRF) with RV (Birn et al., 2008; Chen et al., 2020) in the NKI dataset, while in the HRV-ER dataset, end-tidal CO_2_ was modeled with the an end-tidal CO_2_ response function (Golestani et al. 2015). The peaks and troughs in these impulse response functions represent the characteristic temporal dynamics of physiological signal propagation into the BOLD signal. The use of flexible basis sets can capture differences in how fluctuations in HR, RV, and end-tidal CO_2_ drive BOLD signal changes across different regions and by age.

Regions such as the insula, OFC, ACC, and basal ganglia showed significant age differences in BOLD variance explained by physiological signals, which is noteworthy because these areas have been shown to exhibit reduced cerebral blood flow (CBF) with age (Lu et al., 2011). This suggests that diminished vascular reactivity or neurovascular coupling (linked with central autonomic activity) in these regions could contribute to the observed reductions in physiological-BOLD coupling in older adults. Additionally, BOLD variance explained in white matter by physiological signals was significantly greater in younger adults. Modulation of vascular tone accompanying low-frequency fluctuations in systemic physiology may form a major component of white-matter fMRI signals, particularly in periventricular white matter (Özbay et al., 2018, 2019). The observed age-related reduction of white-matter effects may align with previous studies implicating reduced structural integrity of white matter tracts due to arterial stiffening (Badji et al., 2019; Han et al., 2021). Arterial stiffening can compromise microvascular function, impairing oxygen delivery and potentially reducing the responsiveness of white matter to autonomic signals.

The finding of significant differences in the ventricles is also particularly intriguing. Younger adults showed greater fMRI signal variance explained by physiological signals in the ventricles, which might relate to cerebrospinal fluid (CSF) dynamics. Previous work suggests that negative cerebrovascular reactivity (CVR) in brain ventricles during fMRI reflects a dilation of ventricular vessels, decreasing the relative proportion of CSF per unit volume (Thomas et al., 2013). If these ventricular vessels are less responsive in older adults, it could explain the reduced variance in fMRI signals explained by physiological signals in these regions. Moreover, changes in breathing have been shown to directly alter both white-matter and CSF signals through mechanisms such as sympathetically mediated effects on gray matter vascular tone (Picchioni et al., 2022). Our observations suggest that these global vascular dynamics may also be impaired in aging.

In the HRV-ER dataset, although older adults showed slightly higher percent variance of the BOLD signal explained by CO_2_ in regions such as the occipital cortex—a somewhat counterintuitive pattern given other age-related differences—cross-correlation analyses also revealed that older adults had stronger BOLD-CO_2_ correlations in the ventricles at particular lags. By contrast, in the larger NKI dataset, younger adults typically demonstrated higher PVE by RV throughout much of the brain. One possibility is that the Golestani et al. (2015) basis set for end-tidal CO_2_, derived from younger adults, may not fully capture older adults’ altered vascular responses. However, these findings may also suggest that mechanisms other than arterial CO_2_ concentrations—such as respiratory-correlated heart rate changes (De Meersman, 1993; Masi et al., 2007) and sympathetic nervous system activity on cerebrovasculature (Balasubramanian et al., 2019; Mather, 2024; Rim et al., 2022)—could drive age differences in respiration-related brain hemodynamics in the NKI dataset. Because RV and CO_2_ were collected in separate datasets with different sample sizes, it remains unclear whether the observed discrepancies reflect genuine physiological differences or are partly influenced by sampling variation, reduced power, or different measurement modalities (RV vs. CO_2_). Future studies that measure RV and CO_2_ concurrently will be essential to replicate and further investigate these effects.

It is important to note that the PVE by physiological signals in the BOLD signal is a fractional measure rather than an absolute one. Thus, an observed decrease in PVE for older adults does not necessarily reflect the raw amplitude of physiological responses alone; instead, it can arise from changes in both the numerator (physiologically and neuronally driven variance) and the denominator (total BOLD variance). Therefore, one potential counterargument to our interpretation would be that higher levels of resting-state BOLD variance could account for lower PVE by physiological signals in the BOLD signal in older adults. However, this scenario is unlikely as it is well documented that resting-state BOLD variance declines with age (Garrett et al., 2010; Kumral et al., 2020; Millar et al., 2020; Tsvetanov et al. 2021). Alternatively, if neuronal variability increases in older adults or if neuronal stimuli generate proportionally more BOLD variance than physiological fluctuations, the fraction of variance explained by physiological signals could appear lower—even if absolute physiological responses remain unchanged. Yet empirical evidence generally suggests that aging diminishes neuronal hemodynamic responses of the brain (Fabiani et al., 2014; Ward et. al., 2015; West et al., 2019). For example, older adults show reduced hemodynamic response function amplitude during a visual-auditory task (West et al., 2019). In sum, the most likely explanation for our observed decrease in physiological PVE with age is that vascular and structural changes attenuate the absolute BOLD response to physiological fluctuations (Badji et al., 2019; Han et al., 2021; Hoiland et al., 2019; Lu et al., 2011; Reeve et al., 2024; Rucka et al., 2015). Future studies that simultaneously measure neuronal and physiological components of BOLD variability, along with direct indices of vessel compliance or vascular tone, will help clarify these mechanisms.

### Section 4.4: Time Lag Dynamics and Cerebrovascular Response

In both younger and older adults, the HR-BOLD and RV-BOLD cross-correlations demonstrated canonical patterns, resembling the shapes of the cardiac response function (CRF) and respiratory response function (RRF), respectively. These patterns are characterized by an initial positive peak followed by a decline, transitioning to negative correlations at later lags (*Supplemental Figure 2*). The temporal profiles and regional distributions of these correlations align with findings from previous studies (Birn et al., 2008; Chen et al., 2020; Gold et al., 2024), which similarly reported early positive correlations transitioning to negative correlations as a hallmark of BOLD-physiological coupling. Across both NKI and HRV-ER datasets, the mean HR-BOLD cross-correlation was higher in younger adults in areas that are highly vascularized, including the orbitofrontal cortex (OFC), basal ganglia, ventricles, and white matter tracts supplied by major arteries such as the anterior cerebral artery (ACA), middle cerebral artery (MCA), and posterior cerebral artery (PCA). While the HRV-ER differences did not survive TFCE correction, the dataset showed similar trends as those in the NKI dataset. Since the HRV-ER dataset has a much smaller sample size, it is possible that, with more statistical power, these differences would have reached significance.

The CO_2_-BOLD correlations in our study provide further insight into the role of cerebrovascular response and vascular propagation in age-related changes in neurovascular dynamics. First, *Figure 6* reveals a notable consistency between the CO_2_-BOLD responses in the HRV-ER dataset and the (negative deflection of the) RV-BOLD responses in the NKI data, a dynamic relationship that may be expected from prior reports with concurrent RV and end-tidal CO_2_ monitoring (Chang & Glover, 2009b). In the HRV-ER dataset, younger adults exhibited faster CO_2_-BOLD cross-correlation peaks compared to older adults, which could translate to a more rapid cerebrovascular response to changes in metabolic demands (Hoiland et al., 2019; Lu et al., 2011). Similarly, in the NKI dataset, younger adults showed an earlier onset of the dip in RV-BOLD cross-correlations, which may mirror this faster cerebrovascular response, suggesting a consistent age-related pattern across datasets. Cerebrovascular response, defined as the ability of blood vessels to dilate in reaction to increases in CO_2_, is crucial for maintaining oxygen delivery and cerebral perfusion. In younger adults, this response is more synchronized and efficient, leading to quicker adjustments in blood flow (Hoiland et al., 2019; Lu et al., 2011). In contrast, older adults exhibited delayed CO_2_-BOLD correlation peaks, which may reflect a slower cerebrovascular response and cerebral blood flow due to age-related arterial stiffening and reduced vessel compliance (Badji et al., 2019; Hoiland et al., 2019; Lu et al., 2011; Reeve et al., 2024; Rucka et al., 2015). These changes impair the dynamic ability of the vascular system to transmit blood flow signals efficiently, delaying the response to CO_2_ fluctuations.

### Section 4.5: HRV Biofeedback and Physiological Modulation

Our findings regarding HRV biofeedback highlight intriguing age- and condition-dependent effects. In the HRV-ER dataset, the Osc-condition (in which participants’ instructed goal was to reduce HR oscillations) led to an increase in BOLD variance explained by HR and CO_2_ in younger adults, particularly driven by the CO_2_ condition. This result suggests that diminishing physiological oscillations may amplify the brain’s sensitivity to changes in physiological signals in younger adults, potentially by altering the dynamics of cerebrovascular response. However, it remains unclear whether increasing the brain’s sensitivity to physiological signals above its normal baseline in younger adults is inherently beneficial. While heightened coupling may reflect more efficient neurovascular interactions, excessive sensitivity could also lead to greater susceptibility to physiological noise or dysregulation (Bekar et al., 2012).

In contrast, the Osc+ intervention (which aimed to enhance HR oscillations) in older adults resulted in more negative HR-BOLD cross-correlations at later time lags, closely resembling the patterns typically observed in younger adults. This suggests that increasing HR oscillations through biofeedback may partially restore age-related declines in the cerebrovascular response. Such a shift toward a more <youth-like= physiological response profile may reflect enhanced autonomic flexibility and improved vascular responsiveness (Deschodt-Arsac et al. 2018; Fournié et al. 2021; Mohapatra et al. 2024) – perhaps even with implications for neural activity (Mather and Thayer, 2018; Bright et al., 2020).

Indeed, as proposed by Mather and Thayer (2018), high-amplitude oscillations in heart rate may enhance functional connectivity within emotion-regulation networks—especially in medial prefrontal areas sensitive to physiologically driven oscillatory input. Additional evidence of the potential neural benefits of higher-amplitude heart rate oscillations in older adults comes from Jung et al. (2024), who found that changes in resting HRV significantly mediated reductions in negative emotion in the group instructed to increase HR oscillations (Osc+). Thus, when older adults successfully enhanced their cardiac dynamics, they also experienced improved emotional well-being—an effect directly tied to changes in autonomic functioning (Mather and Thayer 2018). Our observations are also in line with recent work examining other aspects of the HRV-ER dataset, which demonstrate multiple benefits associated with increasing HR oscillations. For instance, Yoo et al. (2022) reported that daily practice to augment HR oscillations (Osc+) led to increased cortical volume in the left orbitofrontal cortex (OFC) for both younger and older adults, suggesting a positive impact on key prefrontal regulatory regions. Beyond emotion regulation, Min et al. (2023) show that Osc+ training decreased plasma Alzheimer’s disease (AD)-related biomarkers (plasma Aβ40 and Aβ42), whereas the Osc− condition actually increased these biomarkers. This suggests a link between autonomic regulation and pathways implicated in AD pathophysiology.

Taken together, these convergent findings demonstrate that interventions designed to boost heart rate oscillations can partially restore age-related declines in autonomic and vascular responsiveness. Our results also suggest that HR-BOLD coupling may serve as a biomarker of autonomic health in older adults, aligning with broader evidence that links increasing vagal tone to enhanced emotional well-being and potentially to neuroprotective benefits.

### Section 4.6: Role of Physiological State

Interestingly, the differences in magnitude of HR-BOLD correlations between older and younger adults (*Figure 4, 5*) suggest potential parallels with the vigilance-related fMRI-autonomic covariance patterns observed by Gold et al., (2024). Their study highlighted that fMRI-autonomic covariance differs across baseline vigilance states, with stronger variance in fMRI explained by physiological signals during low vigilance, which is associated with reduced arousal and parasympathetic dominance. In contrast, high vigilance states, characterized by heightened norepinephrine levels and sympathetic dominance, were associated with weaker fMRI-autonomic coupling. Older adults in our study, with their observed patterns of delayed HR-BOLD and RV-BOLD correlations, may resemble this higher vigilance state. Sympathetic dominance in older adults (Mather, 2024) could explain these temporal differences – particularly as the older-adult cohorts in our study exhibited reductions in high-frequency heart rate variability and RMSSD, compared to the younger adults.

In addition to the analysis described in previous sections, we sought to clarify whether the age-related changes in BOLD-physiological differences were arising due to age-induced changes in the coupling of physiological signals to the BOLD signal, or just age-related changes in physiological signals themselves (Jandackova et al., 2016; Reardon & Malik, 1996; Voss et al., 2012). There is also evidence of sex differences in HRV (Koenig and Thayer, 2018) as well as sex differences in vascular aging (Ji et al. 2018), which may have influenced our results. To isolate the effect of aging on BOLD-physiological coupling as the underlying mechanism explaining our findings, we replicated our PVE and cross correlation analysis while controlling for variables such as sex, average heart rate, LF HRV, HF HRV, breathing rate, average end-tidal CO2 (for the HRV-ER dataset), and standard deviation of RV (in the NKI dataset). Across both datasets, we observed that controlling for these covariates had no effect on the age-related differences in BOLD-HR, BOLD-RV, or BOLD-CO_2_ cross-correlations (*Supplemental Figure 4*). Additionally, in the HRV-ER dataset, adjusting for average HR, LF HRV, and HF HRV had no impact on the Osc+ mediated difference in HR-BOLD cross-correlation among older adults (*Supplemental Figure 5*). However, when controlling for average CO_2_ and breathing rate, the Bonferroni-corrected significant difference in CO_2_-BOLD cross-correlation observed in younger adults before and after the Osc-intervention (*Figure 7d*) was no longer present, though uncorrected significant cross-correlation differences remained (*Supplemental Figure 5)*.

In the NKI dataset, controlling for gender, average HR, LF and HF HRV, standard deviation RV, and breathing rate seemed to preserve most of the significant age-related differences in BOLD PVE by HR and RV in the gray matter and periventricular white matter, although there appears to a reduction in significant voxels in white matter (*Supplemental Figure 6*). This seems to suggest that some of the white matter PVE differences outside of the periventricular area may be attributable to group differences in baseline physiology, whereas the core age effect in the gray matter reflects genuine alterations in physiological-BOLD coupling. Interestingly, in the HRV-ER dataset, controlling for gender, LF and HF HRV, average HR, and average end-tidal CO2 increases the number of voxels in which younger adults’ BOLD PVE by HR and end-tidal CO_2_ is significantly greater than older adults, particularly in gray matter regions, such as the orbitofrontal cortex, anterior cingulate cortex, and insula (*Supplemental Figure 7*). However, additionally controlling for breathing rate removes all significant voxels (*Supplemental Figure 7*), suggesting that breathing rate is tightly linked to the observed coupling differences in this cohort. Taken together, these findings underscore the complexity of interpreting physiological-BOLD coupling in aging research, as different physiological covariates can either reveal or mask significant group differences, depending on how they overlap with core age-related changes in cerebrovascular responsiveness.

### Section 4.7: Limitations and Future Directions

Several limitations should be noted. First, the canonical response functions (CRF/RRF) and the Golestani et al. (2015) basis functions used to model the BOLD response to heart rate, respiration, and CO_2_ were originally derived from younger adults. It is possible that these impulse responses do not fully capture the altered vascular dynamics characteristic of older adults (e.g., increased arterial stiffness or prolonged hemodynamic latency), potentially underestimating the percent variance explained in this group. While our model-free cross-correlation analyses largely corroborate the presence of age-related differences, developing age-specific physiological response functions would be a valuable direction for future work.

Second, the temporal resolution for all of our cross-correlation analyses is inherently limited by the image TR, and future studies with shorter TR may be able to better capture the temporal dynamics of physiological-BOLD coupling. Third, the relatively small sample size for post-intervention scans in the HRV-ER dataset limits statistical power, making it more challenging to detect subtle effects and interactions. Fourth, participants’ adherence to the daily HRV biofeedback protocols may have varied, and differences in training engagement or compliance could influence the observed intervention effects. Fifth, structural differences between younger and older adult brains (e.g., age-related atrophy or enlarged ventricles) may not be fully accounted for by standard preprocessing pipelines, potentially introducing systematic biases in image registration and alignment. Future work could use age-specific templates or advanced normalization methods to better accommodate these structural differences. Sixth, as we focused here on older adults in the range of 50-85 years old, it is possible that the specific age-range may impact the results of old versus young group comparisons. Further, differences in imaging parameters and preprocessing pipelines could introduce technical variability in characterizing relationships between fMRI and peripheral physiological signals. Future studies employing multi-modal imaging (e.g., near-infrared spectroscopy or arterial spin labeling) and longitudinal follow-ups could provide a more comprehensive view of how autonomic regulation interventions influence cerebrovascular function and healthy brain aging over time.

## Data and Code Availability

All source code and instructions for replicating the results in this paper can be found at: https://github.com/richardwsong/physio-bold-aging

Preprocessed Physiological and fMRI data for the HRV-ER dataset is accessible here: https://vanderbilt.box.com/s/ff388e0q9yzx2qkab273q01p9ieo2c2a

The Enhanced Nathan Kline Institute -Rockland Sample dataset is publicly available at the following repository: http://fcon_1000.projects.nitrc.org/indi/enhanced/. The HRV-ER dataset is publicly available at https://openneuro.org/datasets/ds003823.

## Author Contributions

RS: Data preprocessing, Conceptualization, Formal analysis, Methodology, Visualization, Writing–original draft, Writing–review and editing. JM: Data curation, Writing–Review and Editing, Methodology. SW, KRO, SG: Data preprocessing, Writing–review and editing, Methodology. RY: Data preprocessing. HJY, KN: Data curation. JC: Investigation, Writing– review and editing. MM, CC: Conceptualization, Investigation, Methodology, Writing–original draft, Writing–review and editing, Project Administration, Funding Acquisition

## Ethics Statement

For the HRV-ER dataset, The University of Southern California Institutional Review Board approved the study. All participants provided written, informed consent prior to participation and received monetary compensation for their participation. The NKI dataset was openly provided and the study was approved by the Institutional Review Board at the NKI.

## Supplementary Material

**Supplemental Figure 1.**
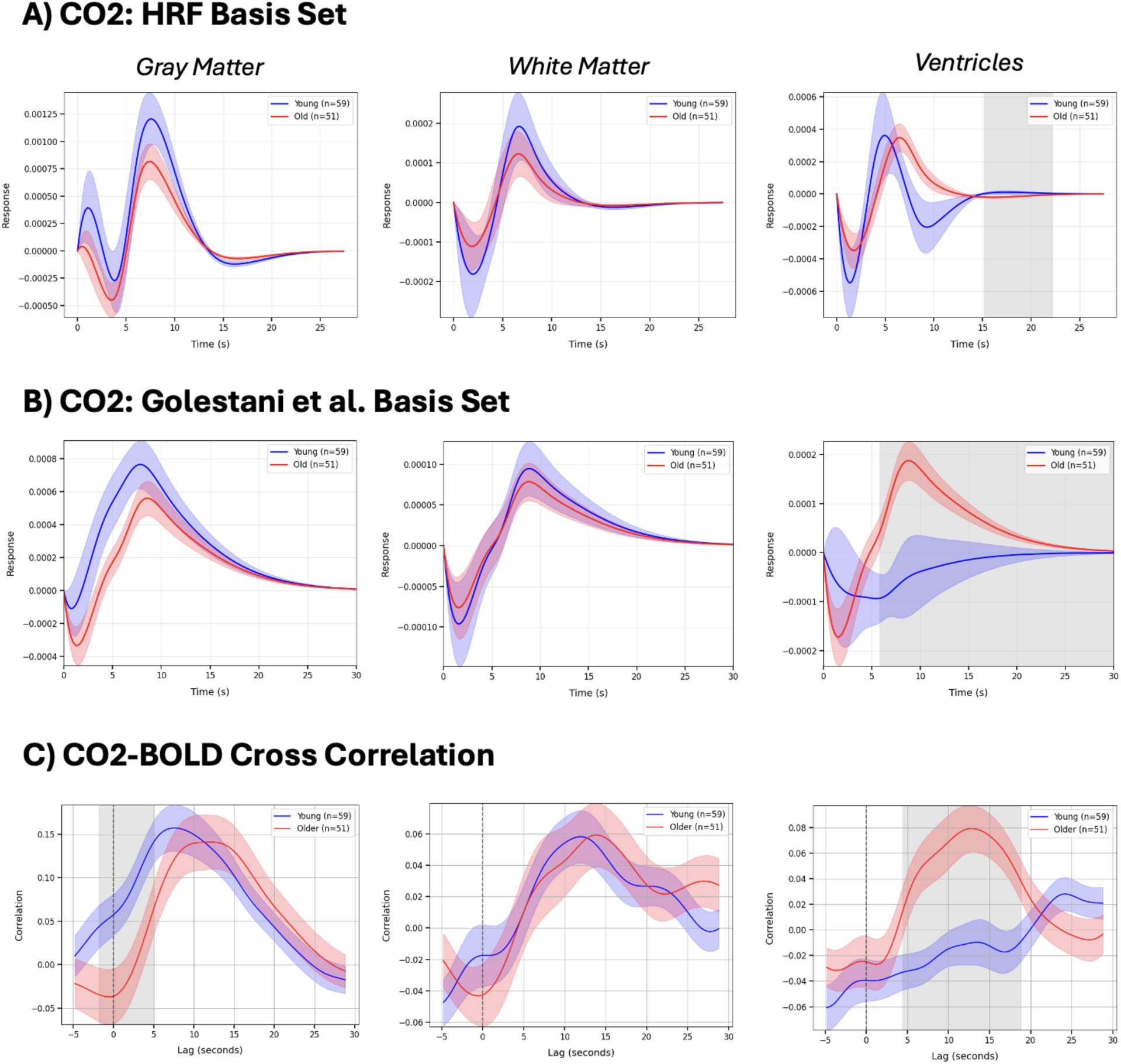
Modeling end-tidal CO2 response with 2 candidate basis sets A) Canonical HRF and B) Golestani et al. (2015) Basis sets were derived by taking a weighted sum of the respective beta values with each basis function after least squares fitting between convolved end-tidal CO_2_ regressors and the BOLD signal averaged across gray matter, white matter, or ventricles. In the HRV-ER data, both CO_2_ and BOLD signals were unsampled to TR = 0.2 seconds before least squares fitting. C) CO_2_-BOLD cross-correlation is shown for reference. Lags/time-points where the response or correlation is statistically significant (*p*<0.05) between older and younger adults are shaded in gray.

**Supplemental Figure 2.**
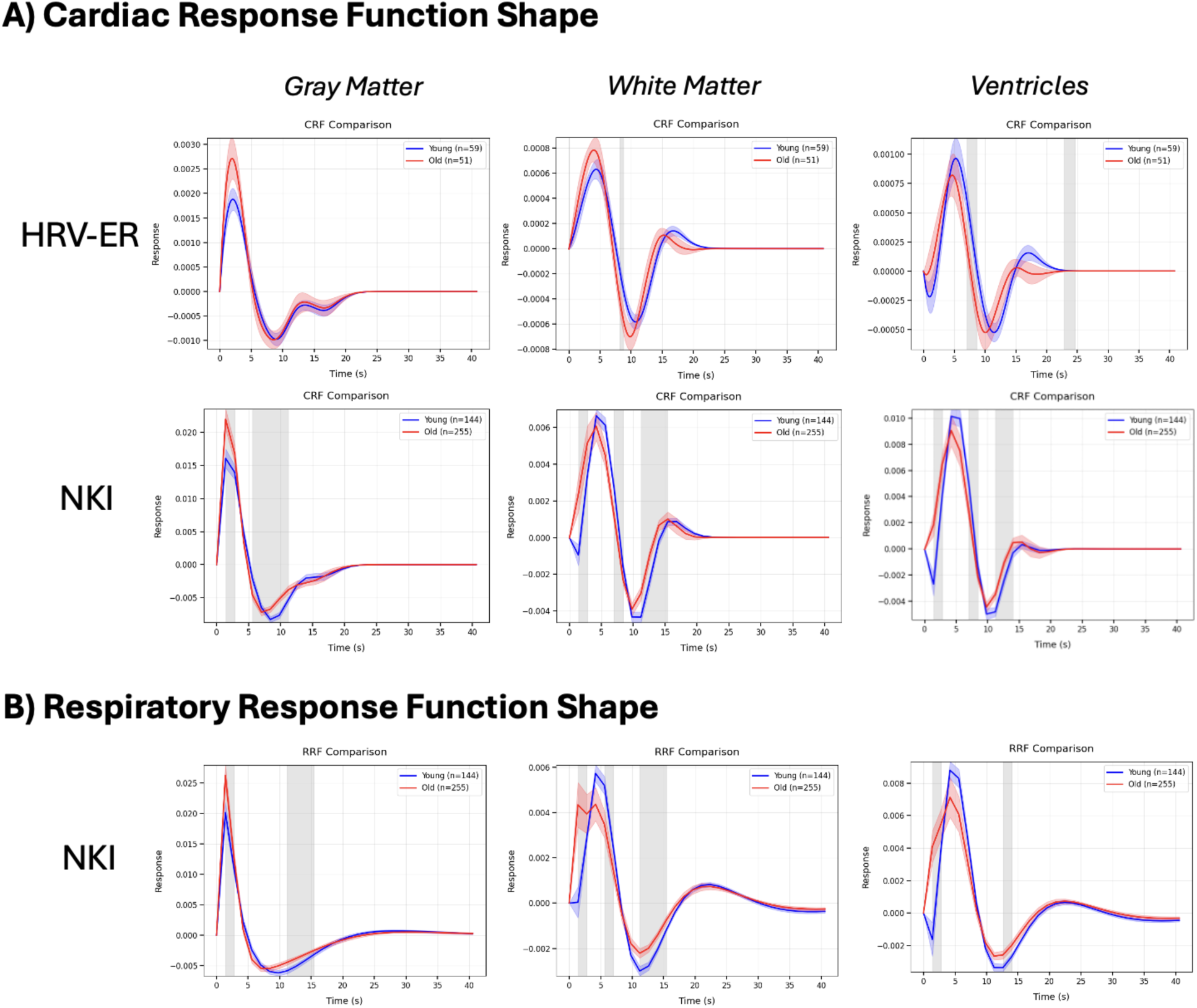
Modeling HR- and RV-induced BOLD response using A) Cardiac Response Function (CRF) and Respiratory Response Function (RRF). Basis sets were derived by taking a weighted sum of the respective beta values with each basis function after least squares fitting between convolved HR/RV regressors and the BOLD signal averaged across gray matter, white matter, or ventricles. In the HRV-ER data, both HR and BOLD signals were unsampled to TR = 0.2 seconds before least squares fitting. Time-points where the response or correlation is statistically significant (p<0.05) between older and younger adults are shaded in gray.

**Supplemental Figure 3.**
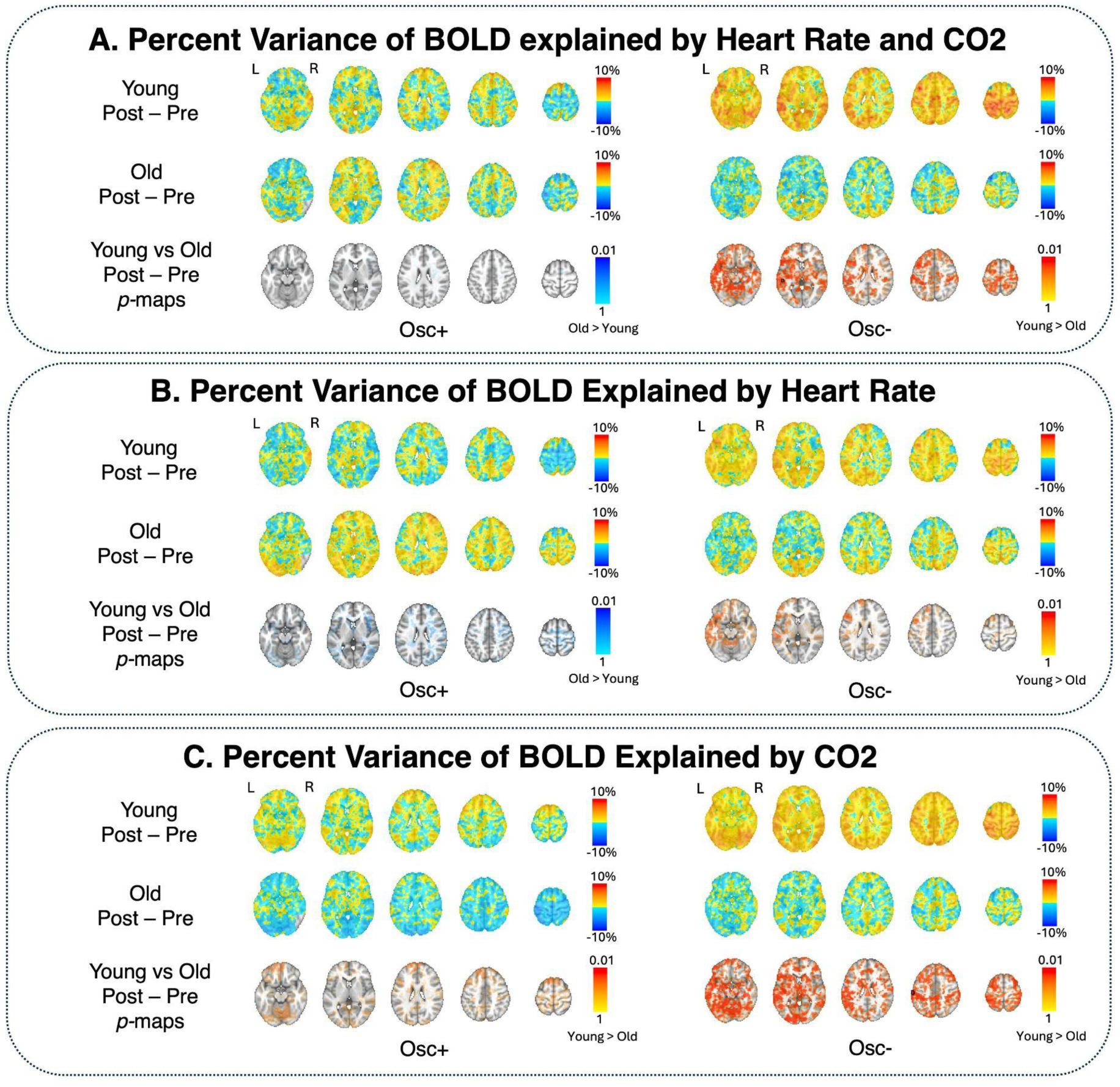
Difference in percent variance of BOLD explained by A) HR and CO2, B) HR, and C) CO2 between pre and post 5 week HRV-biofeedback intervention in the HRV-ER dataset. In each panel, group averages for young adults and older adults are shown in the first two rows, and p-maps comparing the two age groups are shown in the third row. Significant voxels at *p <* 0.05 (TFCE-corrected) are outlined in black, and alpha-fading was used to depict subthreshold voxels.

**Supplemental Figure 4.**
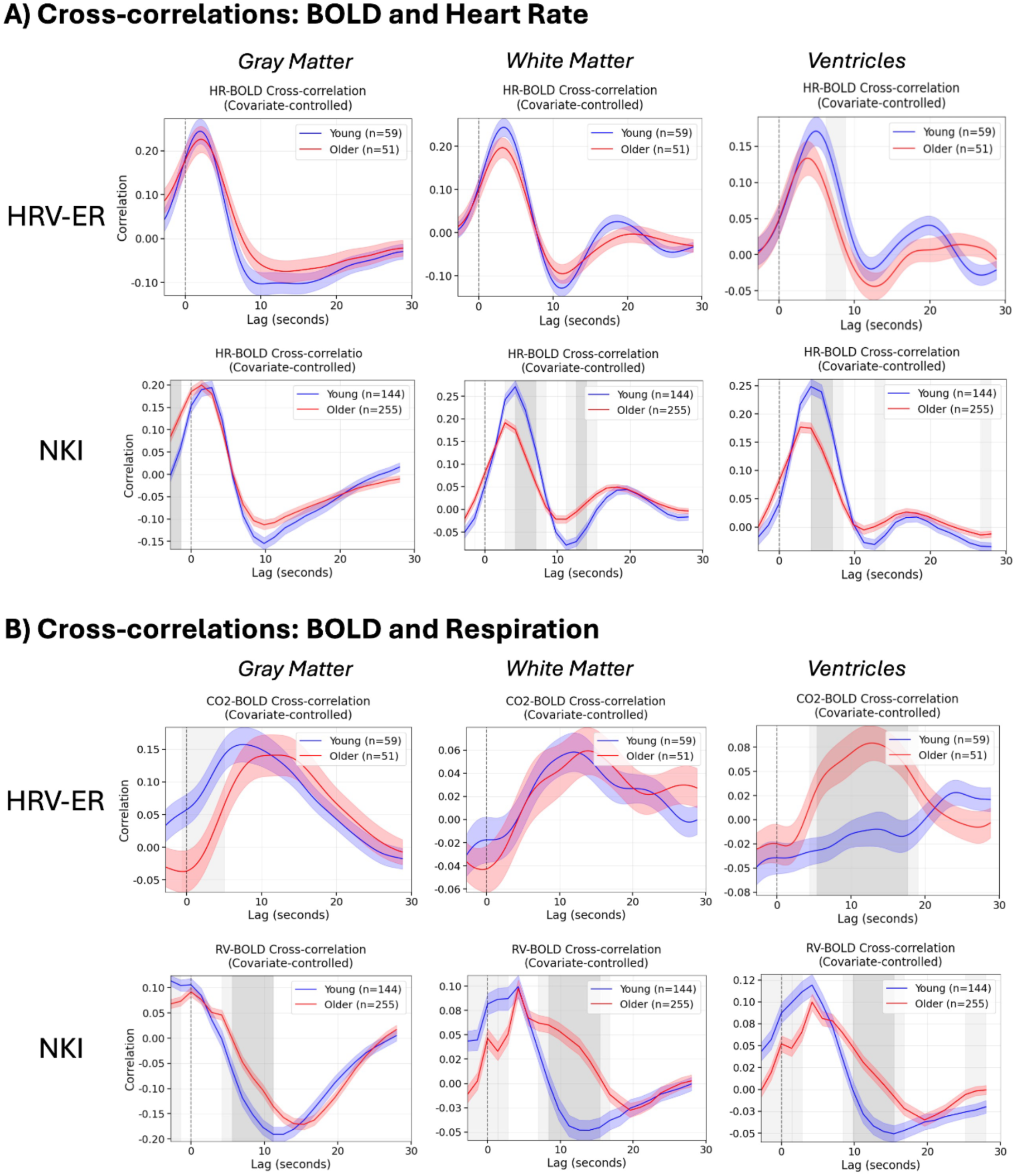
Cross-correlations between the BOLD signal and A) heart rate and B) respiration (i.e., CO_2_ for HRV-ER, RV for NKI), averaged across three tissue types: gray matter, white matter, and ventricles. Lags where the cross correlations between older and younger adults were statistically significant (*p*<0.05) after Bonferroni correction are plotted in dark gray, and lags where the cross correlation passed a *p*<0.05 uncorrected threshold are plotted in light gray. HR-BOLD cross-correlation significance tests were corrected for average heart rate, LF HRV, HF HRV, and RMSSD. CO_2_-BOLD cross-correlations were corrected for average CO_2_ and breathing rate, and BOLD-RV cross correlations were corrected for average RV and breathing rate.

**Supplemental Figure 5.**
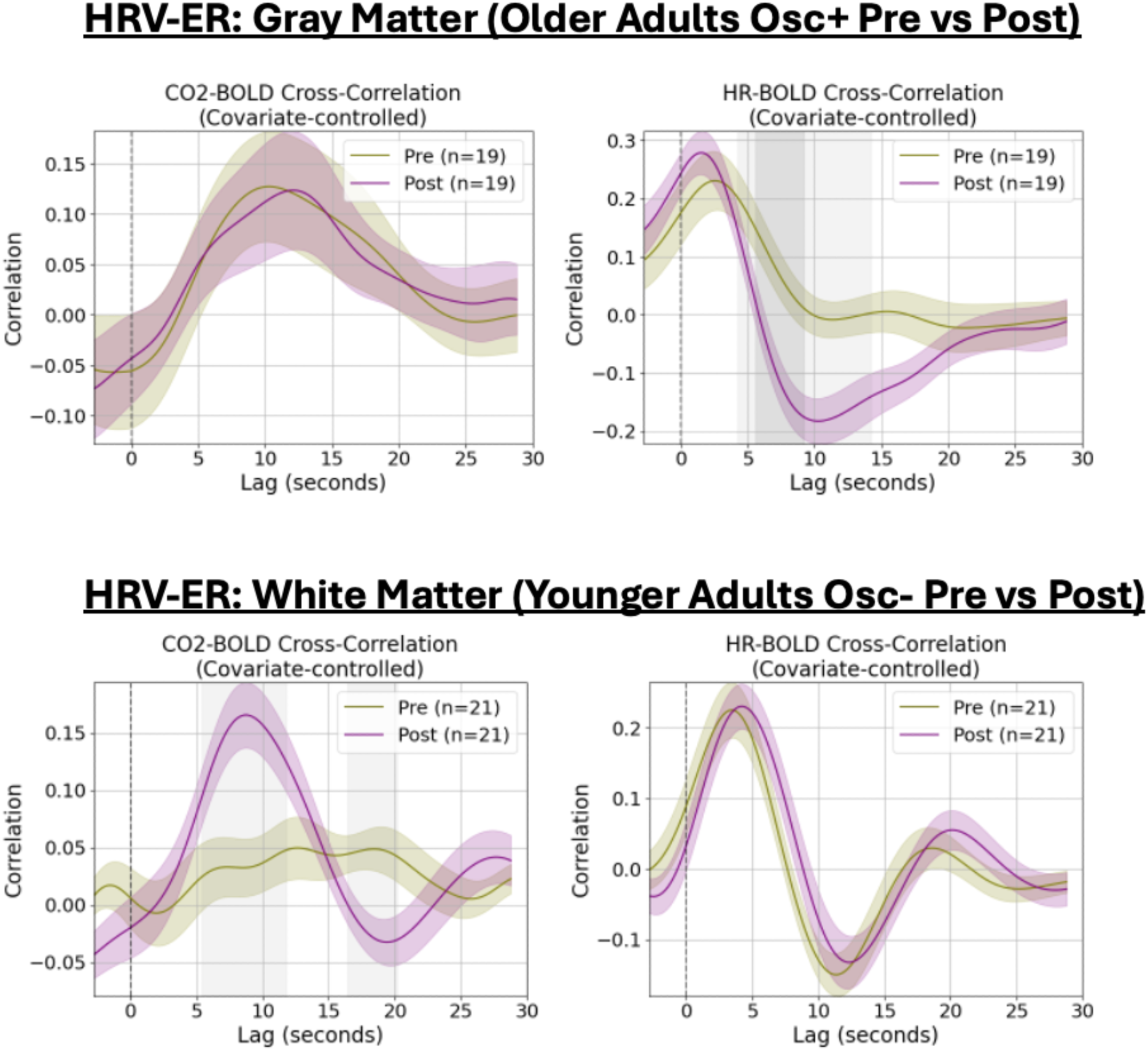
: A) Cross-correlations between CO2 and HR with BOLD signal in the gray matter for older adults before and after the Osc+ intervention. B) Cross-correlations between CO2 and HR with BOLD signal in the white matter for younger adults before and after the Osc-intervention. Lags where the cross correlations between pre and post intervention were statistically significant (p<0.05) after Bonferroni correction are plotted in dark gray, and lags where the difference passed a p<0.05 uncorrected threshold are shown in light gray. Both analyses initially showed significant Bonferroni-corrected differences in Figure 7 of the main text. HR-BOLD cross-correlation significance tests were corrected for average heart rate, LF HRV, HF HRV, and RMSSD. CO_2_-BOLD cross-correlations were corrected for average CO_2_ and breathing rate.

**Supplemental Figure 6.**
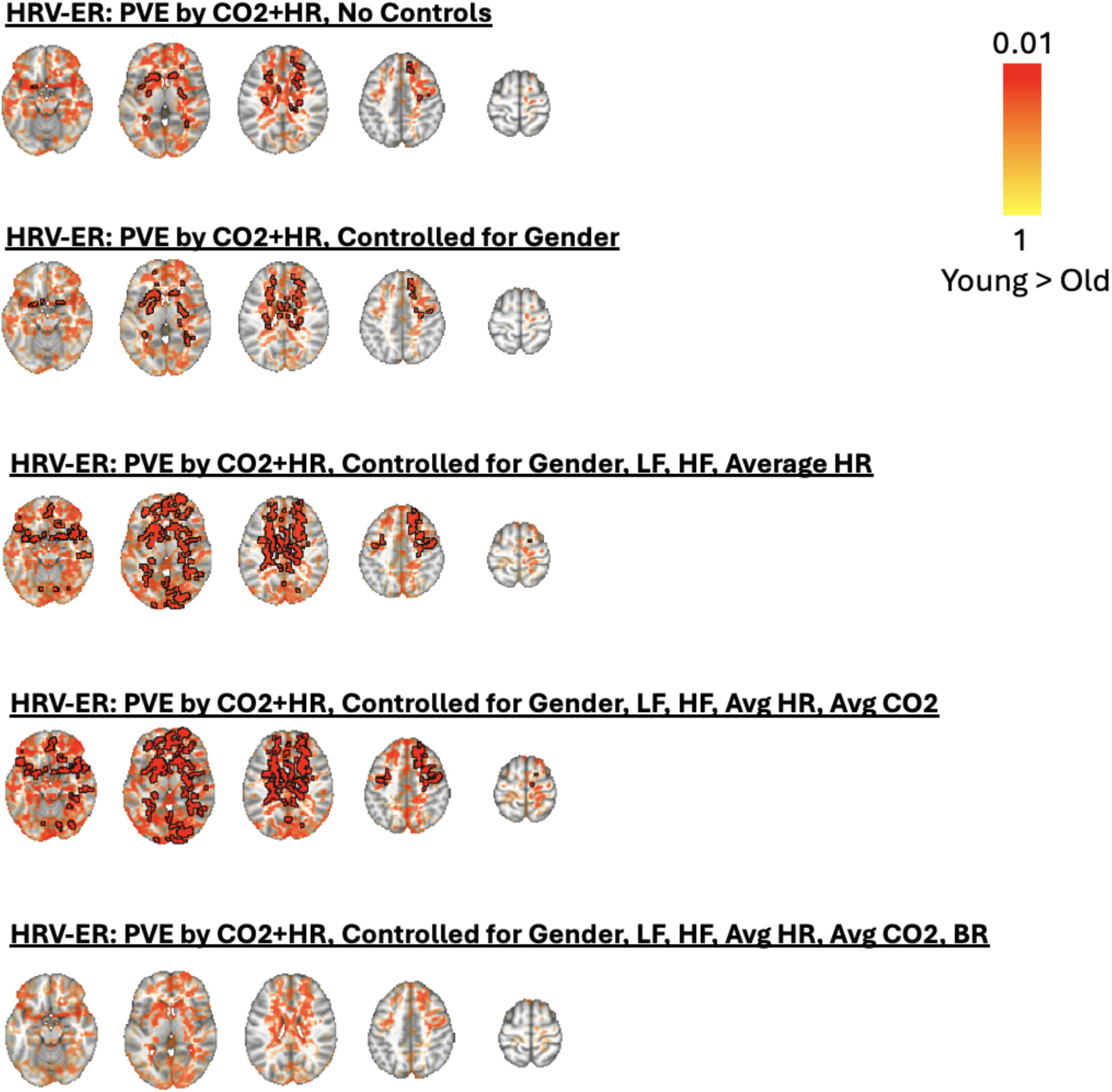
Percent variance of BOLD signal explained by HR and CO_2_ across all voxels in the HRV-ER dataset. Voxels in which percent variance explained in younger adults was statistically significantly greater than older adults (*p* < 0.05 TFCE-corrected) are outlined in black, and alpha-fading was used to highlight sub-threshold voxels.

**Supplemental Figure 7.**
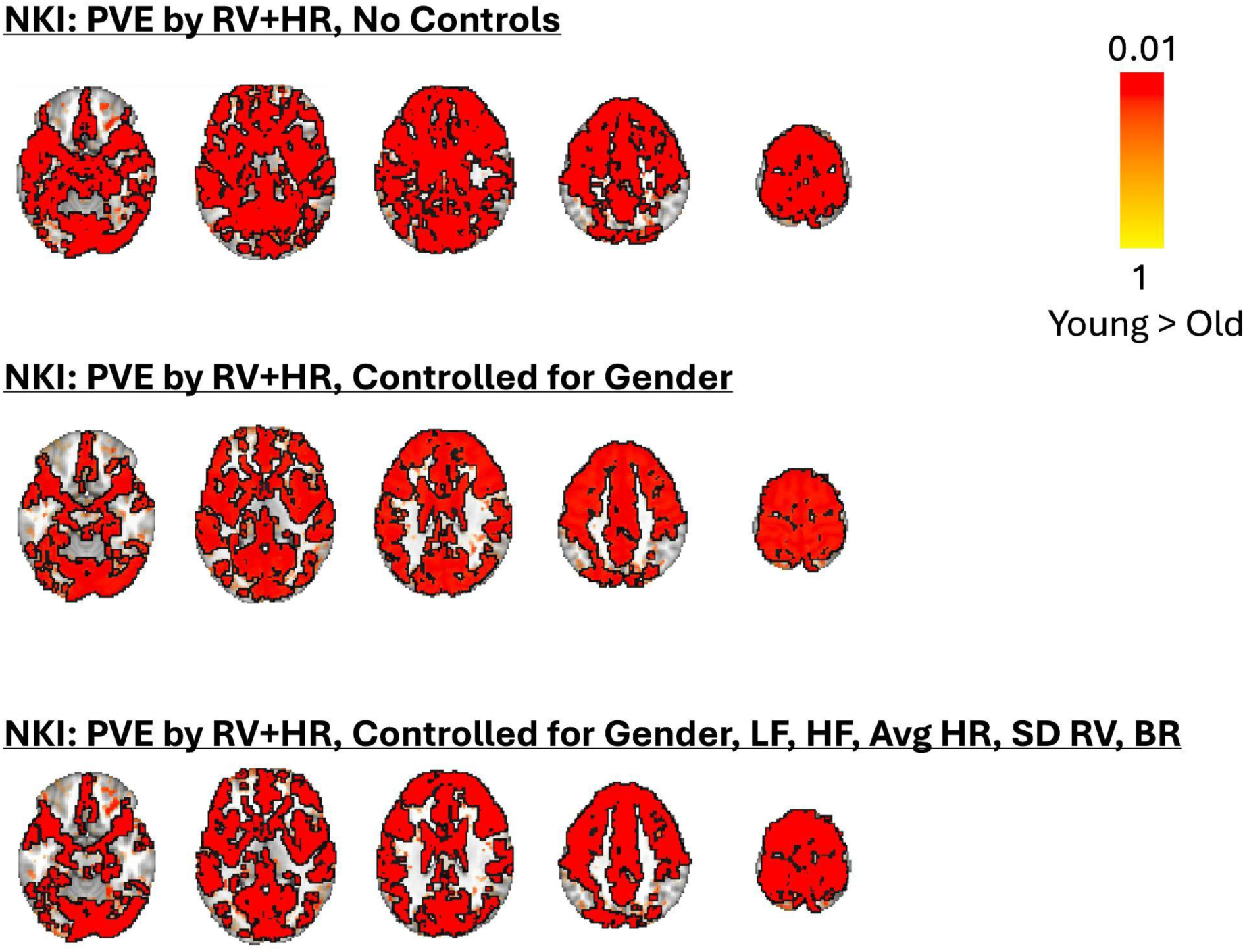
Percent variance of BOLD signal explained by HR and RV across all voxels in the NKI dataset. Voxels in which percent variance explained in younger adults was statistically significantly greater than older adults (*p* < 0.05 TFCE-corrected) are outlined in black, and alpha-fading was used to highlight sub-threshold voxels.

